# Small RNA bidirectional crosstalk during the interaction between wheat and *Zymoseptoria tritici*

**DOI:** 10.1101/501593

**Authors:** Xin Ma, Nicolas Bologna, Javier Palma-Guerrero

## Abstract

Cross-kingdom RNAi has been shown to play important roles during plant pathogen interactions. But this cross-kingdom RNAi was still unexplored in the *wheat-Zymoseptoria tritici* pathosystem. Here we performed a detailed analysis of the sRNA bidirectional crosstalk between wheat and *Z. tritici*. Using a combination of sRNA-seq and mRNA-seq we were able to identify known and novel sRNAs and study their expression and their action on putative targets in both wheat and *Z. tritici*. We predicted the target genes of all the sRNAs in either wheat or *Z. tritici* transcriptome and used degradome analysis to validate the cleavage of these gene transcripts. We could not find any clear evidence of a cross-kingdom RNAi in this pathosystem. We also found that the fungal sRNA enrichment was lower *in planta* than during *in vitro* growth, probably due to the lower expression of the only Dicer gene of the fungus during plant infection. However, we found a downregulation of specific wheat sRNAs during the fungal infection, leading to a boost expression of wheat defense related genes, which may be enhancing the plant defense ability against the pathogen. Additionally, the fungal infection also induced sRNAs regulating the expression of specific wheat genes, including auxin related genes, as an immune response. These results confirm the role of sRNAs in the regulation of wheat defenses during *Z. tritici* infection. Our findings contribute to improve our understanding of the interactions between wheat and *Z. tritici*.

## Introduction

Small RNAs (sRNAs) are short non-coding RNAs (approximately 20 to 30 nucleotides in length) that play important roles in the regulation of gene expression and genome stability in eukaryotic organisms (Moazed, 2009). The three major classes of sRNAs are microRNAs (miRNAs), small interfering RNAs (siRNAs) and piwi-associated RNAs (piRNAs) (Carthew and Sontheimer, 2009; Moazed, 2009). While miRNA and siRNA are highly extended among plants, piRNAs have so far only been found in animal cells and are necessary in the development of germ cells by silencing transposons or other genetic elements (Vagin et al., 2006; Siomi et al., 2011).

In plants, miRNAs (~21nt) are produced from imperfect fold-back structures of self-complementary non-coding transcripts, originated from endogenous loci, and they regulate target mRNAs by Post-Transcriptional Gene Silencing (PTGS). In contrast, siRNAs derive from double strand RNA (dsRNA) with near-perfect complementarity, and they can be both of *endogenous origin*, e.g. inverted repeat (IR) and dsRNA synthesized by endogenous RNA-dependent RNA polymerases (RDRs) (Allen et al., 2005; Felippes and Weigel, 2009; Fei et al., 2013; Creasey et al., 2014), or *exogenous origin*, e.g. viral intermolecular replication intermediates or intramolecular secondary structures. siRNAs guide either PTGS (mediated by 21/22nt siRNAs) or RNA-directed DNA methylation (RdDM) often resulting in transcriptional gene silencing (TGS; mediated by 24nt siRNAs) (Matzke and Mosher, 2014; Borges and Martienssen, 2015). While each sRNA pathways possess specific characteristics in terms of biogenesis and accessory proteins, the basic mechanism of RNA silencing share few consensus biochemical steps: (*i*) initiation by dsRNA, (*ii*) dsRNA processing to generate the mature sRNAs by Dicer (or Dicer-like, DCL) proteins, and (*iii*) sRNA incorporation into Argonaute (AGO) proteins to interact with target mRNA or DNA in order to execute their silencing functions (Liu et al., 2004; Hutvagner and Simard, 2008; Bologna and Voinnet, 2014).

sRNAs are involved in plant development, reproduction, genome reprogramming, and contribute to the phenotypic plasticity of plants (Borges and Martienssen, 2015). Besides, plant sRNAs also mediate the response to abiotic stresses, such as water stress, sulfate stress, phosphate starvation, and cold stress (Sunkar and Zhu, 2004; Sunkar et al., 2007; Park et al., 2010; Guleria et al., 2011; Kamthan et al., 2015). Plants as well employ RNAi as defense strategies against biotic stress (Ruiz-Ferrer and Voinnet, 2009; Zhang et al., 2011; Weiberg et al., 2014). In Arabidopsis, miR393 is induced by the bacteria elicitor flg22 promoting the pathogen-associated molecular patterns (PAMPs) triggered immunity (PTI) (Navarro et al., 2008). MiR393 suppresses the auxin signaling by targeting auxin receptors TIR1 and three related F-box proteins, this results in increased plant resistance against the pathogen (Achard et al., 2007; Chen et al., 2007; Grant and Jones, 2009). MiR482, which can target numerous NBS-LRR genes in tomato, reduces its levels in response to bacterial pathogens and some viruses, resulting in increased target NBS-LRR transcripts in the immune system (Shivaprasad et al., 2012). This miRNA mediated NB-LRR regulation is considered to be important for plant immunity to pathogens (Zhai et al., 2011; Li et al., 2012; Shivaprasad et al., 2012; Fei et al., 2013).

In fungi, RNAi is an ancient host defense mechanism against viruses and transposons invasion (Chang et al., 2012; Nicolas and Ruiz-Vazquez, 2013; Torres-Martinez and Ruiz-Vazquez, 2017). However, no miRNAs, but only miRNA-like RNAs (milRNAs) have been identified in fungi (Lee et al., 2010; Goodwin et al., 2011), which share similarities with conventional miRNAs, but are synthesized by different mechanisms (Lee et al., 2010). The first RNAi process in fungi, called quelling, was found in *Neurospora crassa*. Quelling is triggered by repetitive transgenes during mitotic growth, and it is essential for suppressing transposons replication (Romano and Macino, 1992; Nolan et al., 2005). QiRNAs, a class of Quelling-Deficient 2 (QDE2) interacting sRNAs, are specifically induced when *Neurospora* is treated with a DNA-damaging agent (Lee et al., 2009). Additionally, sRNAs are also required for fungal growth and pathogenesis in *Magnaporthe oryzae* (Raman et al., 2017).

RNAi has recently been reported to play important roles in plant and pathogen interactions. In host-induced gene silencing (HIGS), accumulation of double-stranded or antisense RNAs in barley and wheat can target and silence genes in the fungal pathogen *Blumeria graminis* as an effective defense strategy (Nowara et al., 2010). This cross-kingdom RNAi has also been shown to happen in a natural way. A fungal pathogen, *Botrytis cinerea*, can hijack host plant defenses by using small RNAs that bind to the plant AGO1 and silence plant defense related genes (Weiberg et al., 2013). A novel milRNA has been identified in *Puccinia striiformis f. sp.tritici* (Pst) that silences a wheat pathogenesis-related 2 gene and influences the plant immunity (Wang et al., 2017). Plants can also use the same RNAi mechanisms to defend against pathogenic fungi, e.g. cotton plants can export their miRNAs into the fungal pathogen *Verticillium dahliae*, and inhibit the expression of virulence genes during the infection cycle (Wang et al., 2016; Zhang et al., 2016). In addition, a recent study has shown that *Arabidopsis* can send sRNAs in extracellular vesicles to fungal pathogens, revealing how sRNAs travel from hosts to fungal pathogens (Cai et al., 2018).

However, no previous studies have reported the roles of sRNAs in the pathosystem of wheat and the fungal pathogen *Zymoseptoria tritici*, even though this fungus is the causal agent of the septoria tritici blotch, the most damaging disease on wheat in Europe (Hardwick et al., 2001; O’Driscoll et al., 2014) and an important disease of wheat worldwide. After penetrating wheat leaves through stomata, *Z. tritici* starts the complex infection cycle characterized by a long period of asymptomatic growth known as a latent period that can up to two weeks (Sanchez-Vallet et al., 2015). This latent period is followed by a rapid switch to necrotrophic growth in which a large number of plant cells are killed in a two to three days period, and it is accompanied by a rapid accumulation of fungal biomass inside the plants (Rudd et al., 2015). So far, only one effector gene, LysM, has been identified in *Z. tritici*, which is required for virulence and can suppress PTI during the infection (Marshall et al., 2011). Therefore, the mechanisms underlying *Z. tritici* pathogenicity still remain largely unknown. In recent studies we have analyzed the transcriptomes of both wheat and *Z. tritici* during the infection (Palma-Guerrero et al., 2017; Ma et al., 2018). These studies revealed the mutual transcriptomic responses between wheat and *Z. tritici* during the interactions, which, together with other transcriptomics studies in this pathogen, shows that a whole genome wide reprogramming of gene expressions happens in both plant and fungus (Rudd et al., 2015; Palma-Guerrero et al., 2017; Ma et al., 2018). But the role of sRNAs and the RNAi in the wheat-Z. *tritici* pathosystem remains largely unknown. Besides, no previous study described this natural bidirectional RNAi between plant and pathogen at the same time in the same samples, which could provide a detailed scenario of the communications between plant and pathogens.

Here we analyzed the small RNAs produced during wheat infection by *Z. tritici* at key stages of the disease cycle to identify both the *Z. tritici* sRNAs and the wheat sRNAs and their degraded targets. We found fungal sRNAs induced during infection and predicted their targeted genes in the wheat transcriptome. But cleavage of these predicted targets could not be confirmed by degradome analysis. Additionally, the expression of the fungal dicer gene was downregulated and barely expressed *in planta*. At the same time, we found that wheat could downregulate sRNA expressions and lead to an increase defense ability against *Z. tritici*. Additionally, wheat sRNAs involved in the regulation of auxin signaling were also induced in response to *Z. tritici*. Thus, our results suggest that the cross-kingdom RNAi in wheat and *Z. tritici* pathosystem does not lead do detectable mRNA degradation by PTGS during infection. However, RNAi can be used as an immune response in plants cells.

## Materials and Methods

### Samples collection and RNA extractions

The total RNAs from leaf samples of wheat cultivar drifter infected with the *Z. tritici* 3D7 strain at 7, 12 and 14dpi (days post infection) were obtained in a previous study from our group (Palma-Guerrero et al., 2017). The RNAs from uninfected mock samples at 7, 12 and 14dpi that serve as control were obtained in this study, by using the control samples generated in the same previous study (Palma-Guerrero et al., 2017). The RNA samples of the fungus growing *in vitro* were generated in a different previous study (Francisco et al., 2018). Three replicates per sample with the highest RNA quality according to Bioanalyzer 2100 (Agilent) were used as biological replicates for library preparation and sequencing.

### Small-RNA-seq analysis

The small-RNA-seq libraries were constructed using the TruSeq small-RNA kit and sequenced by HiSeq4000 by the Functional Genomic Center of Zurich (FGCZ). Reads were trimmed to remove sequencing adapters and keep only insert with length >=15 using Trimmomatic-0.36 (Bolger et al., 2014) with the settings “ILLUMINACLIP:smallRNA-SE.fa:2:30:10 LEADING:2 TRAILING:2 SLIDINGWINDOW:4:15 MINLEN:15”. The adapter sequence in smallRNA-SE.fa is “TGGAATTCTCGGGTGCCAAGG”.

### Wheat sRNA predictions

All the clean reads were collapsed using fastx-ToolKit (Gordon and Hannon, 2010). Wheat tRNAs, rRNAs, and small nucleolar RNAs (snoRNAs) were removed. The sequences of tRNAs, rRNAs and snoRNAs were extracted from Triticum_aestivum.TGACv1.ncrna.fa on Ensembl plant (Kersey et al., 2018).

We used miRDeep-p (Yang and Li, 2011) and ShortStack (Shahid and Axtell, 2014; Johnson et al., 2016) for the wheat miRNA predictions. For miRDeep-p, all the filtered reads were mapped against wheat genome IWGSC RefSeq v1.0 (Appels et al., 2018) using bowtie (Langmead et al., 2009) allowing no mismatches. The mapped reads from all the samples including infected samples and mocks were merged together and collapsed. We used the mapped reads to retrieve miRNAs annotated in IWGSC RefSeq v1.0 annotations (Appels et al., 2018). We used miRDeep-P (Yang and Li, 2011) to predict wheat miRNAs setting the length of candidate precursors as 300nt. A customized Perl script was used to filter the miRNAs based on the new criteria of plant miRNA annotation (Axtell and Meyers, 2018). Basically, 1) both the miRNA and miRNA* are predicted and expressed in the sequencing data; 2) The expression of miRNA is more than 10 reads per million in at least one of the samples; 3) The length of mature miRNAs should be between 20nt and 24nt; 4) The mismatches between miRNA and miRNA* are under 5; 5) The asymmetric mismatches between miRNA and miRNA* are under 3; 6) The length of precursor should be under 300nt.

We as well used ShortStack (Shahid and Axtell, 2014; Johnson et al., 2016) to predict the wheat sRNA loci with the settings “—mismatches 0 –mincov 0.5rpm”. Three replicates of each conditions were merged together for the ShortStack alignment and annotation. Only the sRNAs candidate locus with expressions of at least 10 rpm in at least two replicates of each condition were kept. Then the major siRNAs and miRNAs from every condition were summarized together. We named every unique siRNA and miRNA reads as miRNA-uniq or siRNA, and with ascending order numbers. We combined the annotations of miRNAs from Arabidopsis, rice and wheat from miRBase and the annotations of wheat miRNA from IWGSC together to make a customized database of the known plant miRNAs (Griffiths-Jones, 2004; Griffiths-Jones et al., 2006; Griffiths-Jones et al., 2008; Kozomara and Griffiths-Jones, 2011; 2014). All the unique mature sequences of predicted miRNAs were used to blast against the customized database to detect if the miRNAs were previously annotated (Camacho et al., 2009).

All the sequences of mature sRNAs were used to count sRNAs expression in each sample. We used TMM (Robinson and Oshlack, 2010) to normalize the library sizes and RPM (reads per million) to describe the sRNAs expression. The differentially expressed sRNAs were called by edgeR (Robinson et al., 2010) using log(Fold change) >=1 (or <=-1) and FDR<0.05.

### Fungal sRNA predictions

The fungal tRNAs, rRNAs and snoRNAs were removed from clean reads. The annotations of tRNAs, rRNAs and snoRNAs were extracted from Ensembl fungi (Goodwin et al., 2011; Kersey et al., 2018). The published genome reference of 3D7 was used in the analysis (Plissonneau et al., 2016). Then the filtered reads were used to predict fungal sRNAs with ShortStack, using the same strategy as described above for the plant. The expression of the fungal sRNAs were obtained as described above for the plant sRNAs.

### SRNA target predictions and degradome analysis

The wheat and *Z. tritici* sRNAs upregulated or highly expressed (RPM>=100) at 7dpi or 12dpi were used to predict the target genes in wheat and *Z. tritici* transcriptome using psRNATarget (Dai et al., 2018) Default parameters from Schema V2 (2017 release) were used, but the expectation value was set at 4.5. Only the targeted genes that were significantly downregulated while the sRNAs showed upregulation or were constantly highly expressed were considered as the predicted target genes.

To confirm the cleavage of the predicted target genes, we generated modified PARE libraries with RNAs from the fungus (*in vitro*), infected wheat leaves (7dpi and 12dpi), and mock leaves at the same time points. The samples for the modified PARE libraries were the same ones used for the sRNA-seq libraries. Poly(A)+ RNA was isolated from the RNA samples. Illumina TruSeq sequencing adapter was ligated to the 5’P ends of the degraded mRNA. Next, first-strand cDNA synthesis was performed using a N6 randomized primer. After RNA hydrolysis, the 3’ Illumina TruSeq sequencing adapter was ligated to the 5’ ends of the antisense 1st-strand cDNA fragments. The 5’ cDNA fragments were finally amplified with PCR using the Illumina TruSeq sequencing adapters as primers. The libraries were sequenced on an Illumina Next Seq 500 single end at 1×75bp using a Truseq SBS kit v3-HS (Illumina, Inc. San Diego, CA, USA).

The raw sequencing data were trimmed to remove adapters and trimmed length>=15 using Trimmomatic-0.36 (Bolger et al., 2014) with the settings “ILLUMINACLIP:TruSeq3-SE.fa:2:30:10 LEADING:10 TRAILING:10 SLIDINGWINDOW:4:15 MINLEN:15”. The clean reads were analyzed using CleaveLand4 (Brousse et al., 2014) with the parameters “-c 3 -p 0.05 -r 0.7” for every possible conditions. Only the results with a p value under 0.05 and a minimum free energy (MFE) ratio higher than 0.7 were regarded as significant results. We only considered the degradome peaks in Category 0, 1, 2 and 3. These categories are described in Cleaveland 4.

### mRNA-seq sequencing and analysis

The mRNA-seq data from the wheat samples infected with *Z. tritici* were generated in a previous study (Palma-Guerrero et al., 2017) and they are available at the National Center for Biotechnology Information (NCBI) Short Read Archive (SRA) under the project SRP077418. The new mRNA-seq data from the 3D7 isolate grown *in vitro* and plant mocks at 7dpi, 12dpi were generated in this study using Illumina HiSeq2500 at 2 X 125 bp (NCBI-SRA project number SRP154808). All the raw sequencing reads were trimmed using Trimmomatic-0.36 as described before. The clean reads were aligned against wheat genome and *Z. tritici* genome separately using tophat v2.1.1 (Kim et al., 2013). HTSeq-count v0.6.1 was used to calculate the gene counts (Anders et al., 2015). We used TMM (trimmed mean of M values) implemented in Bioconductor EdgeR v3.12.0 to normalize the library sizes and gene expressions (Robinson et al., 2010; Robinson and Oshlack, 2010). Differentially expressed genes were defined using EdgeR. The Benjamin-Hochberg false discovery rate correction was used to adjust P values based on the exact Fisher test (Robinson et al., 2010). The genes with a log fold change >=1 (or <=-1) and adjusted p value <0.05 were determined as the differentially expressed genes. The GO analysis were performed with TopGO v2.32.0 (Alexa and Rahnenfuhrer, 2016).

## Results

### *Z. tritici* produces sRNAs during wheat infection

To explore the roles of RNAi in the wheat-Z. *tritici* pathosystem, we analyzed sRNA-seq libraries from samples of wheat leaves infected by *Z. tritici* after 7dpi, 12dpi and 14dpi. The corresponding wheat mock leaves at the same points, and *Z. tritici* grown *in vitro*, were used as controls. Three biological replicates were used for each condition. According to the description of the disease cycle (Rudd et al., 2015; Sanchez-Vallet et al., 2015), our dataset covered the latent period (7dpi), transition period (12dpi) and the early necrotrophic period (14dpi) of the 3D7 strain disease cycle (Palma-Guerrero et al., 2017).

Reads with similarity to tRNAs, rRNAs and snoRNAs of wheat and *Z. tritici* were filtered and the remaining reads were aligned against the wheat and the *Z. tritici* genome separately to distinguish the plant and fungal sRNA reads (Table 1). In total, we obtained 0.3 to 1.1 million fungal sRNA reads in the infected samples and 1.8 to 2.8 million fungal sRNA reads in the *in vitro* samples. Between 3.4 to 18.1 million plant sRNA reads were obtained in the infected samples and 1.9 to 6.5 million plant sRNA reads in the mock samples. The number of fungal sRNA reads increased during the infection process while the number of plant sRNA reads decreased, which is consistent with the disease progression on the infected leaves.

**Table 1.** SRNA-seq data summary. sRNA ratios represent the percentage of fungal or plant sRNAs among the total amount of clean reads.

| Conditions | Library ID | Total<br>clean reads | Fungal<br>sRNA reads | Fungal<br>sRNA ratios | Plant<br>sRNA reads | Plant<br>sRNA ratios |
| --- | --- | --- | --- | --- | --- | --- |
| **Infection**<br>**samples** | 3D7-7dpi-2 | 71,764,255 | 339,262 | 0.47% | 18,186,405 | 25.34% |
|  | 3D7-7dpi-8 | 75,365,538 | 385,127 | 0.51% | 17,763,333 | 23.57% |
|  | 3D7-7dpi-10 | 48,440,536 | 306,834 | 0.63% | 9,039,173 | 18.66% |
|  | 3D7-12dpi-2 | 65,969,573 | 460,119 | 0.70% | 12,378,083 | 18.76% |
|  | 3D7-12dpi-5 | 65,138,243 | 653,052 | 1.00% | 10,109,163 | 15.52% |
|  | 3D7-12dpi-6 | 45,790,107 | 317,915 | 0.69% | 10,725,249 | 23.42% |
|  | 3D7-14dpi-4 | 31,090,290 | 901,839 | 2.90% | 3,783,930 | 12.17% |
|  | 3D7-14dpi-5 | 34,372,386 | 577,053 | 1.68% | 5,365,083 | 15.61% |
|  | 3D7-14dpi-10 | 30,549,120 | 1,108,689 | 3.63% | 3,406,298 | 11.15% |
| **Fungal *in vitro*** | 3D7-vitro-1 | 16,547,092 | 2,865,958 | 17.32% | ----- | ----- |
|  | 3D7-vitro-2 | 12,318,911 | 2,191,788 | 17.79% | ----- | ----- |
|  | 3D7-vitro-3 | 9,515,263 | 1,819,328 | 19.12% | ----- | ----- |
| **Plant**<br>**mocks** | mock-7dpi-C1 | 17,166,095 | ----- | ----- | 4,586,381 | 26.72% |
|  | mock-7dpi-C2 | 21,565,083 | ----- | ----- | 5,321,432 | 24.68% |
|  | mock-7dpi-C3 | 24,379,228 | ----- | ----- | 6,568,124 | 26.94% |
|  | mock-12dpi-C1 | 20,048,211 | ----- | ----- | 3,884,980 | 19.38% |
|  | mock-12dpi-C2 | 23,466,121 | ----- | ----- | 4,787,936 | 20.40% |
|  | mock-12dpi-C4 | 14,114,494 | ----- | ----- | 3,065,456 | 21.72% |
|  | mock-14dpi-C1 | 8,993,244 | ----- | ----- | 1,902,557 | 21.16% |
|  | mock-14dpi-C2 | 28,168,941 | ----- | ----- | 6,222,211 | 22.09% |
|  | mock-14dpi-C4 | 11,895,554 | ----- | ----- | 2,884,932 | 24.25% |

Using a customized bioinformatic analysis (see the methods section), we predicted 662 unique fungal sRNAs in total (Table S1) (Shahid and Axtell, 2014; Johnson et al., 2016). We considered all the major RNAs predicted by ShortStack as the fungal sRNAs. The most abundant fungal sRNAs were of 20 and 21 nucleotides (nt) in length (Figure 1A). The fungal sRNAs were originated across the fungal genome, including both core and accessory chromosomes, although most sRNAs came from chromosomes 1 and 3 (Figure 1B). Among the total number of fungal sRNAs, 160 were originated from gene exon regions, 19 were from gene intron regions, 343 were from transposable elements and 212 were from the intergenic regions (Figure 1C).

**Figure 1.**
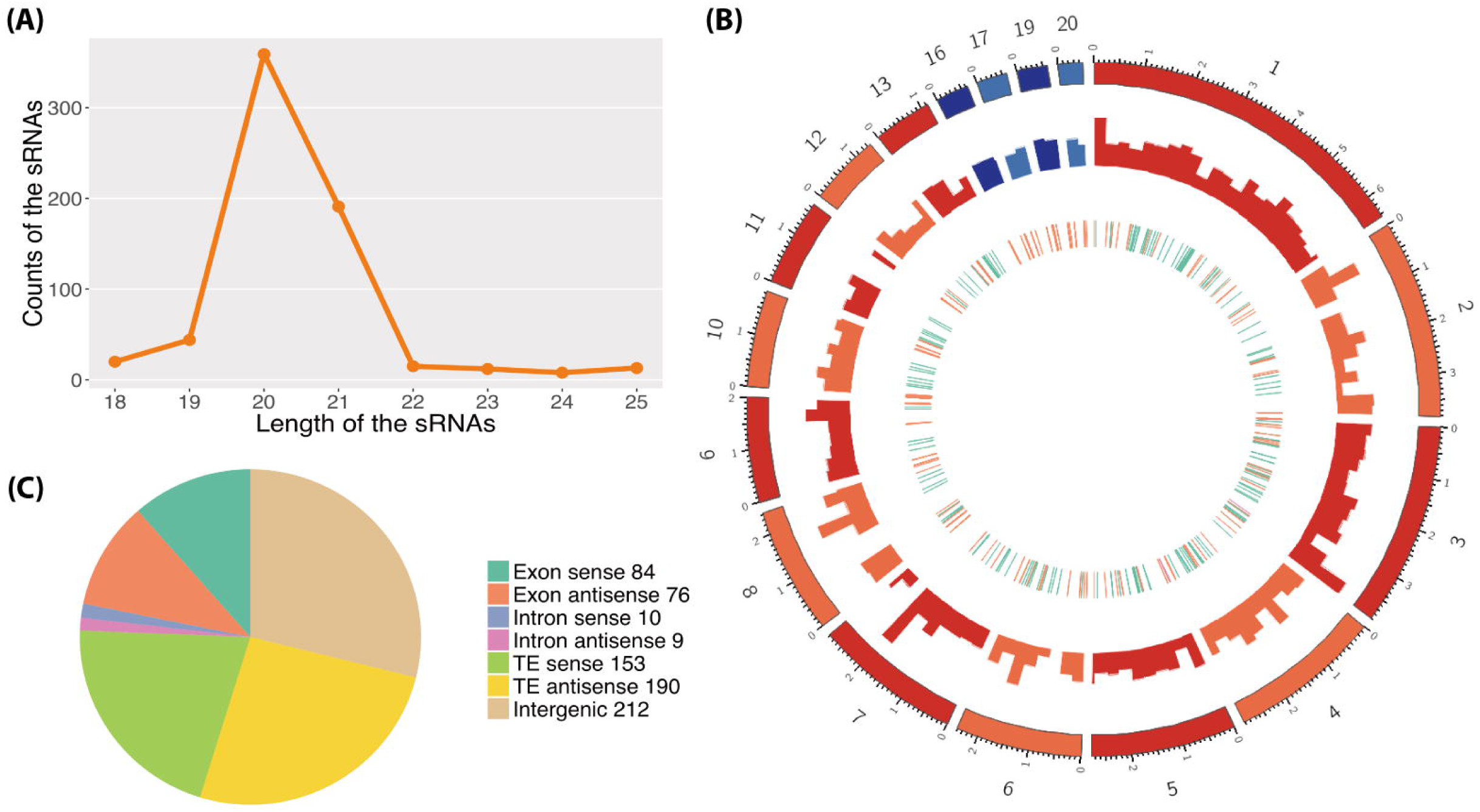
Predictions of the fungal sRNAs. (A) Length distribution of the predicted fungal sRNAs. The x-axis indicates the lengths of the sRNAs and the y-axis indicates the counts of the sRNAs. (B) Distribution of the fungal sRNAs across the fungal genome. The outer layer shows the chromosomes of the fungal genome. The red colors are the core chromosomes and the blue colors are the accessory chromosomes. Accumulation of fungal sRNAs (log2 transformed) are shown in the middle layer. The inner layer shows the genomic elements where fungal sRNAs come from. The green ones reveal the sRNAs comes from gene region, the orange ones from TE and the pink ones from the overlap of gene and TE region. (C) Origin of the fungal sRNAs. The different colors reveal the genomic elements where the fungal sRNAs come from. The sense means the sRNAs come from the same strand with the genomic elements while the antisense means the opposite strand.

### The composition of fungal sRNAs changes during the infection cycle

We analyzed the differentially expressed fungal sRNAs between infected samples and *in vitro* samples to identify the infection induced fungal sRNAs. We detected that 66 fungal sRNAs were significantly upregulated in the infected samples compared to the *in vitro* samples (Figure 2A). These fungal sRNAs were considered as the infection induced fungal sRNAs and therefore they may play important roles during plant infection. The number of induced fungal sRNAs was only 16 at 7dpi, when the fungal pathogen was in the latent period and no symptoms were detected on the leaves. But this number increased to 30 at 12dpi, when the first chlorotic symptoms appeared on the leaves, and 59 at 14dpi, when necrotic lesions are clearly observed on the leaves (Palma-Guerrero et al., 2017). Besides, there were 174 fungal sRNAs significantly downregulated in the infection samples (Figure 2A).

**Figure 2.**
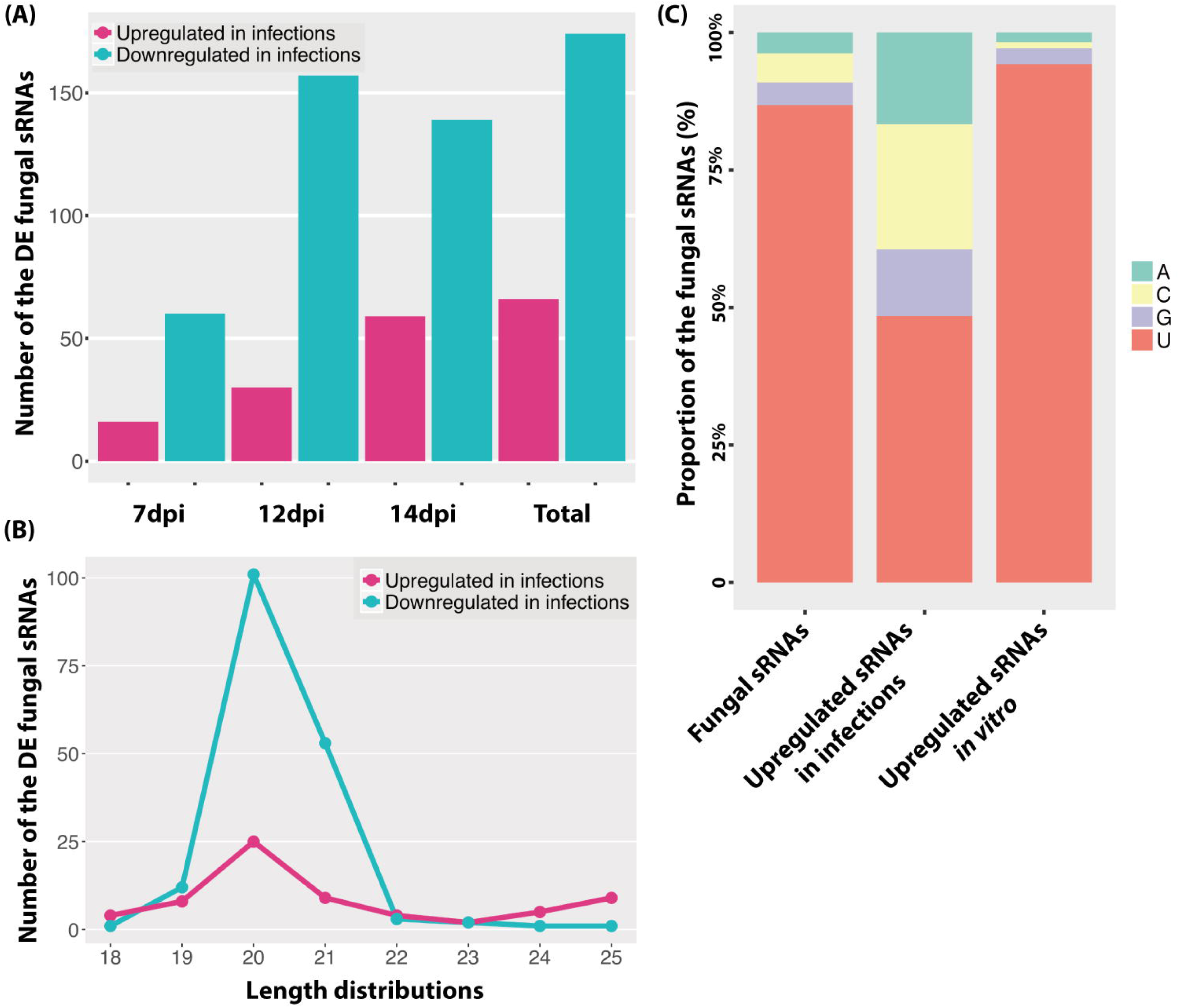
Upregulated sRNAs during the infection cycle. (A) Number of the differentially expressed (DE) fungal sRNAs during the infection cycle. “Total” indicates the total number of DE fungal sRNAs across all the time points. (B) Length distributions of the DE fungal sRNAs. (C) Frequency of nucleotides at 5’ end of the fungal sRNAs.

The length and the nucleotide at 5’ end of the sRNAs is considered to determine to which AGO proteins are they binding, as it has been shown for Arabidopsis, and subsequently the sRNA final function (Mi et al., 2008). In our analysis, the length distribution of the differentially expressed fungal sRNAs between *in vitro* and infected samples were similar being 20/21nt (Figure 1A, 2B). At the same time, most of the fungal sRNAs shared uracil (U) at 5’ end, which have been shown to bind preferentially to AGO1 in Arabidopsis (Figure 2C). Interestingly, the proportion of U at 5’ end changed dramatically, from 87% among all the sRNAs to 48% among the infection induced sRNAs. Correspondingly, 23% of the infection induced fungal sRNAs started with C, binding preferentially to Ago 5 in Arabidopsis, and 16.7% started with A, binding preferentially to Ago2 and Ago3 in Arabidopsis (Figure 2C). This change indicates that there is different fungal RNA accumulation between growth *in vitro* and *in planta*, and that the sRNAs induced during infection could have different functions than the sRNAs produced during *in vitro* growth being part of different AGOs/sRNA complexes.

### *Z. tritici* sRNAs were not detected to cleave wheat transcripts

The cross-kingdom RNAi between plants and fungal pathogens has been shown in different pathosystems (Zhang et al., 2011; Weiberg et al., 2013; Wang et al., 2016; Zhang et al., 2016; Wang et al., 2017). To investigate the presence of cross-kingdom RNAi in the wheat-Z. *tritici* pathosystem, we used the predicted fungal sRNAs to detect target genes in the wheat transcriptome. We found that 33 fungal sRNAs were either highly expressed (above 100 RPM) or upregulated at 7dpi or 12dpi, which were predicted to target wheat genes. Among these targeted wheat genes, 139 were significantly downregulated during the infection (Table S2).

These 139 wheat genes were significantly enriched in chlorophyll related processes and pigment biosynthesis (Figure 3A), which would affect photosynthesis, the leaf color, and the plant defense ability (Menzies et al., 2016). This suggests that fungal sRNAs may regulate plant chlorophyll related genes to reduce the plant defense ability. Interestingly, these targeted wheat genes were enriched in “microtubule-based movement” biological process, which could affect the plant intracellular transport and secretion, and therefore affect the plant response to the pathogen infection (Lee et al., 2012). Besides these, fungal sRNAs were also predicted to target wheat RLK genes and resistance genes, which also suggests that fungal sRNAs were trying to disturb the plant defense system (Table S3).

**Figure 3.**
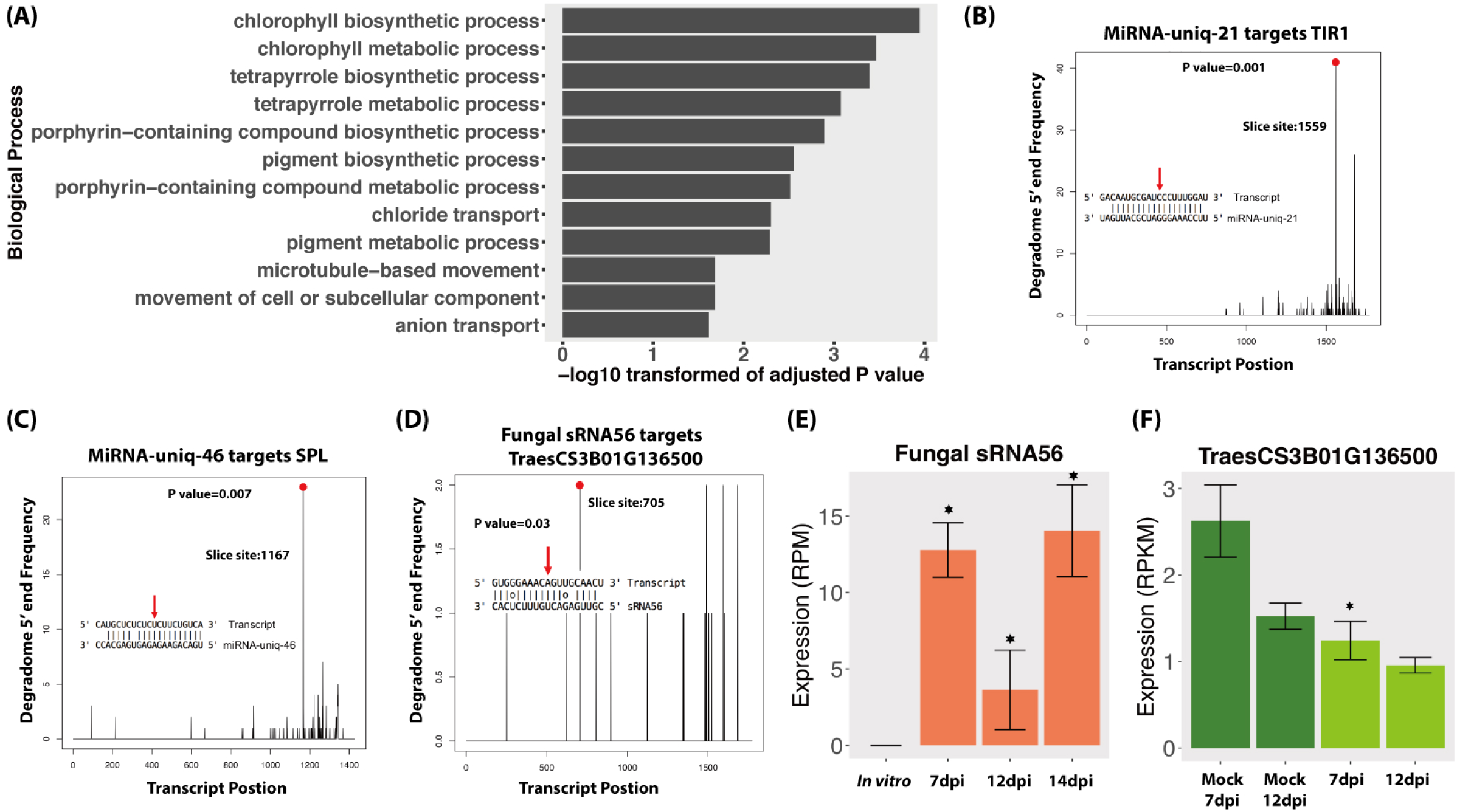
Degradome analysis of the fungal sRNAs. (A) GO enrichment of biological process of wheat genes that were downregulated and predicted to be targeted by fungal induced sRNAs during the infections. (B, C) Positive controls of degradome sequencing results. The x-axis indicates the position of the target transcript and the y-axis indicates the degraded frequency of the target transcript. The red point and red arrow indicate the slice site from the transcript and the sRNA. The P value indicates the significance of the target. (D) Degradome results of fungal sRNA56 targeting plant gene. (E,F) Expressions of fungal sRNA56 and its target gene in plant. The asterisk indicates that the sRNA or the target genes are differentially expressed compared to the *in vitro* or the Mock conditions respectively.

Next, we performed a degradome analysis to validate the cleavage of the candidate targeted genes. Three conditions were used in this analysis: *Z. tritici* grown in vitro, wheat leaves infected by *Z. tritici* (7dpi and 12dpi) and uninfected mock leaves at the same time points. To determine the quality of the degradome sequencing, we first analyzed several plant miRNAs that are already known to target plant genes. In our analysis, these plant miRNAs were predicted to target and cleave plant transcripts as previously reported, which confirmed the good quality of the degradome sequencing. For example, miRNA-uniq-21 (known as miR393) and miRNA-uniq-46 (known as miR156) were correctly found to cleave TIR1 genes and Squamosa promoter-binding-like (SPL) transcription factor genes respectively in our data (Figure 3B,C), as it has been previously reported (Addo-Quaye et al., 2009).

In the degradome analysis, only one wheat gene, TraesCS3B01G136500, was found to be cleaved and downregulated by fungal sRNA56 (Figure 3D). This wheat gene encodes a “Remodeling and spacing factor 1 (RSF-1)” protein, which is known to be involved in DNA double strand break repair in mammals (Pessina & Lowndes, 2014). SRNA56 was induced during infection, but being low expressed (Figure 3E). Additionally, the significantly reduced target, TraesCS3B01G136500, was constantly very low expressed (below 3 RPKM) and the degradome coverage of the cleaved transcripts was also low, which raises doubt about it being a real target (Figure 3D,F).

We also investigated the *RNAi* in *Z. tritici* during the disease cycle. In total, 39 fungal sRNAs were predicted to target and downregulate the fungal own genes during the infection (Table S4). But none of these genes were detected to be cleaved in the degradome sequencing. Finally, we analyzed the expressions of genes involved in RNAi in *Z. tritici*. There is only one dicer gene in the *Z. tritici* genome (g9428 from the 3D7 genome), which is consistent across five different sequenced *Z. tritici* genomes (Goodwin et al., 2011; Plissonneau et al., 2016; Plissonneau et al., 2018). Surprisingly, this dicer gene was significantly downregulated *in planta* compared with *in vitro* (Figure 4A). Additionally, the sRNAs were more abundant *in vitro* than *in planta*, which could be caused by the low expression of the fungal Dicer gene (Figure 4B). There are four Ago genes in the *Z. tritici* genome (g4308, g1839, g10449, and g2094 of the 3D7 genome), among which the Ago gene g2094 was not expressed in any condition tested. We as well did not find any significant upregulation of AGO genes in *Z. tritici* during infection (Figure 4A).

**Figure 4.**
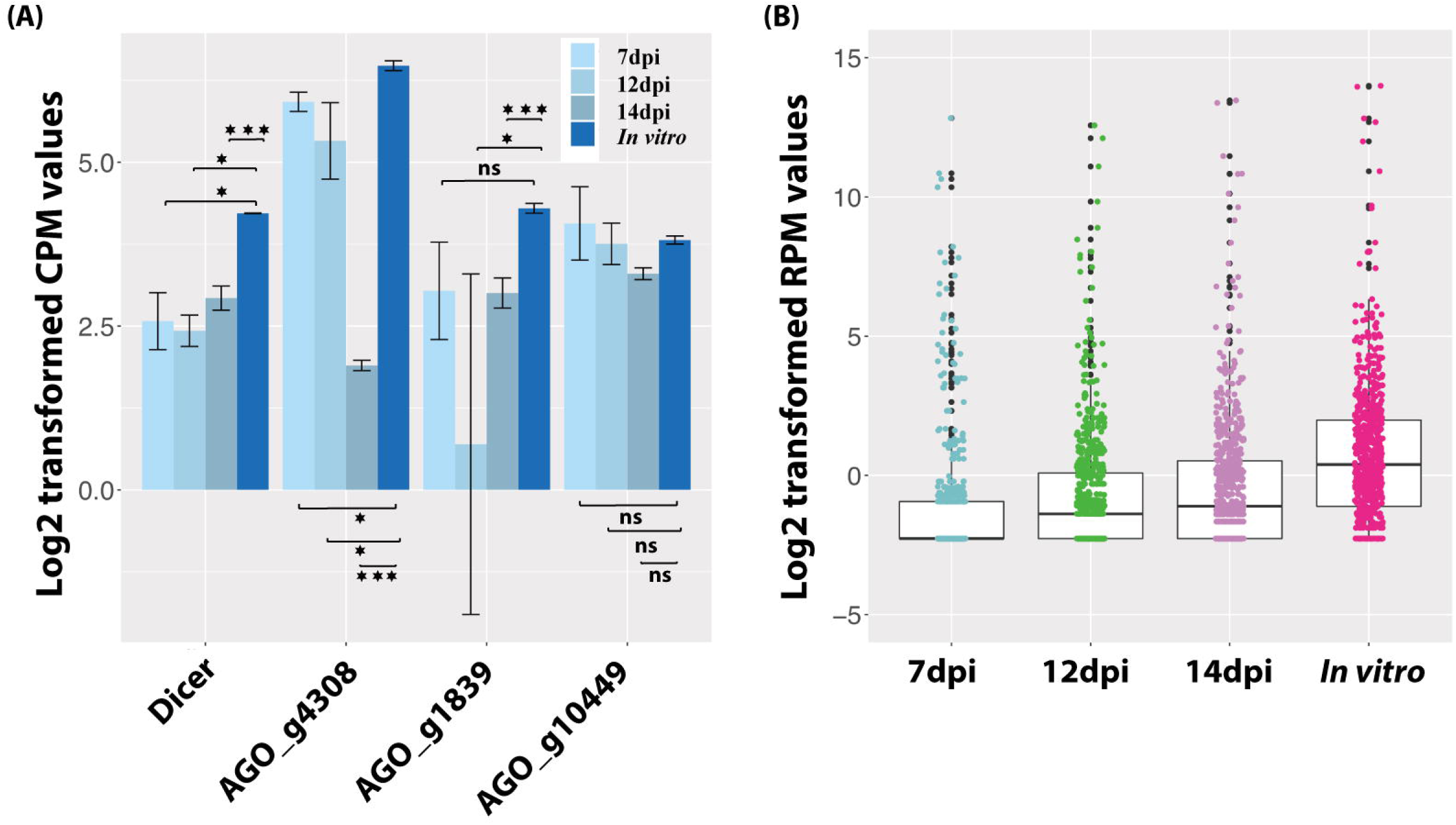
Fungal sRNAs are more abundant *in vitro* than *in planta*. (A) Expression of the fungal Dicer gene and three AGO genes. One asterisk indicates that the adjusted p value is under 0.05 and three asterisks indicates that the p value is under 0.001. ns indicates no significant difference. (B) Average expression of all the fungal sRNAs *in planta* and *in vitro*.

### Wheat regulates sRNAs in response to fungal infections

We used the same dataset to identify the plant sRNAs and to study the roles of plant sRNAs in the disease cycle. In total, we identified 158 wheat miRNAs and 1120 wheat siRNA loci (Table S5). By blasting these predicted miRNAs against the known wheat miRNAs, we found 69 miRNAs that were already annotated in the wheat genome. The other 89 miRNAs were considered as novel wheat miRNAs (Table S5). Most wheat predicted miRNAs were 21nt in length, consistent with previous studies (Figure 5A) (Rajagopalan et al., 2006; Axtell, 2013). The majority of these plant miRNAs start with U at 5’ end (Figure 5B) which should favor binding the plant AGO1 proteins (Mi et al., 2008). Unlike the miRNAs, most of the wheat siRNAs were 24nt in length and started with A (Figure 5A,B). These 24nt siRNAs have been shown to act by RdDM in Arabidopsis (Matzke and Mosher, 2014), being preferentially loaded in AGO4 (Mi et al., 2008).

**Figure 5.**
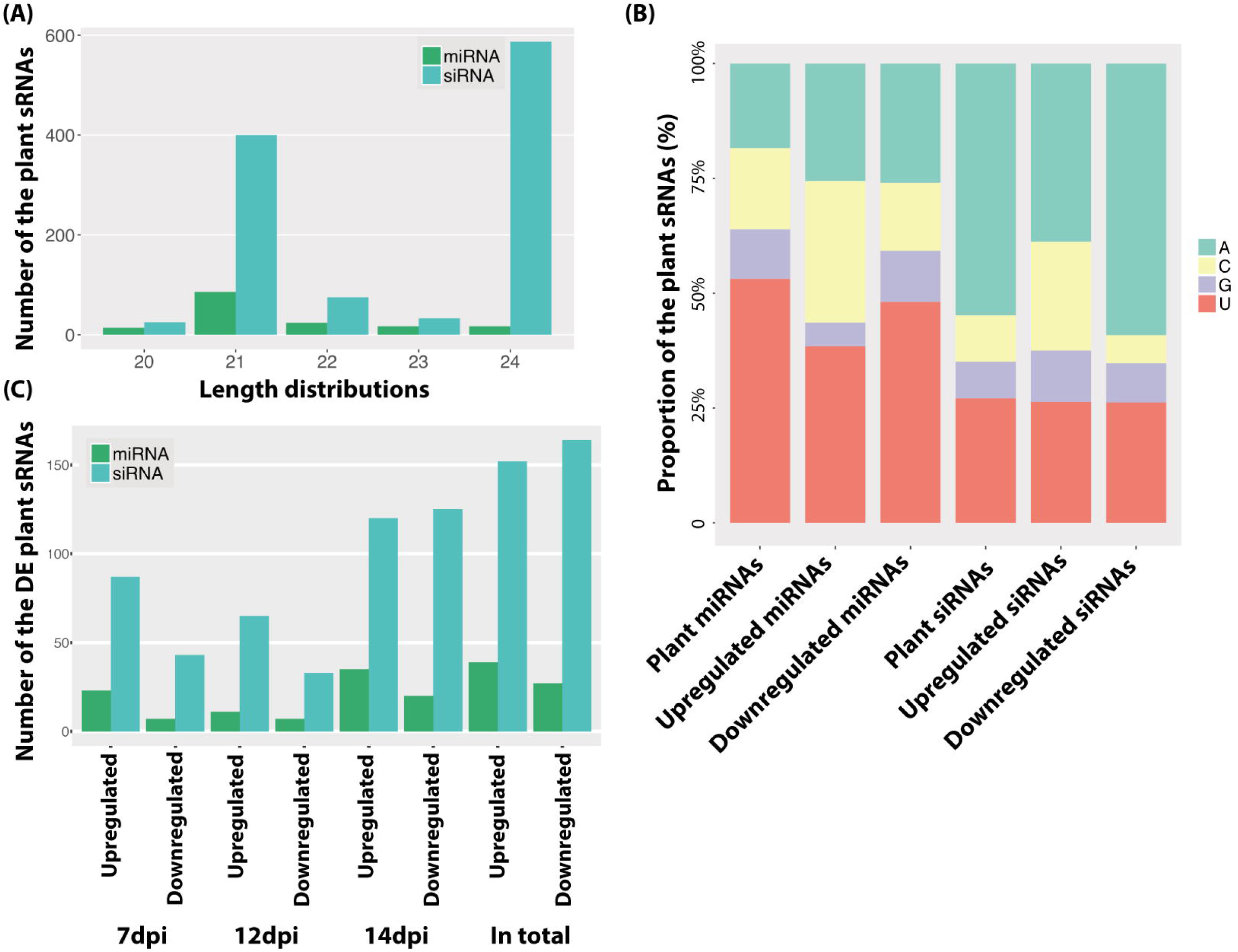
Predicted plant sRNAs. (A) Length distribution of the predicted plant sRNAs. (B) Frequency of nucleotides at 5’ end of plant sRNAs. Different colors indicate different nucleotides. (C) Number of the DE sRNAs between infections and mocks. Upregulated indicates the sRNAs are upregulated in the infections and Downregulated indicates the sRNAs are downregulated in the infections.

Plants can regulate the sRNAs, especially miRNAs, to respond to pathogens, which is thought to be a fast and efficient immune response (Fei et al., 2013). In order to determine if the infection induces variations in sRNAs accumulation, we compared the expression of all the plant sRNAs between infected and mock samples at every time point. Like for the fungus, the infection induced plant sRNAs also exhibited different frequencies of nucleotides at 5’end (Figure 5B), which suggests a different sRNA production during infections. We found 191 plant sRNAs (27 miRNAs and 164 siRNAs) significantly downregulated during the infections (Figure 5C, Table S6). Interestingly, these downregulated plant sRNAs were predicted to target 122 disease resistance genes and 214 genes encoded RLKs in mocks (Table S7). Two RLK genes were detected to be cleaved by siRNA191 (Figure 6).

**Figure 6.**
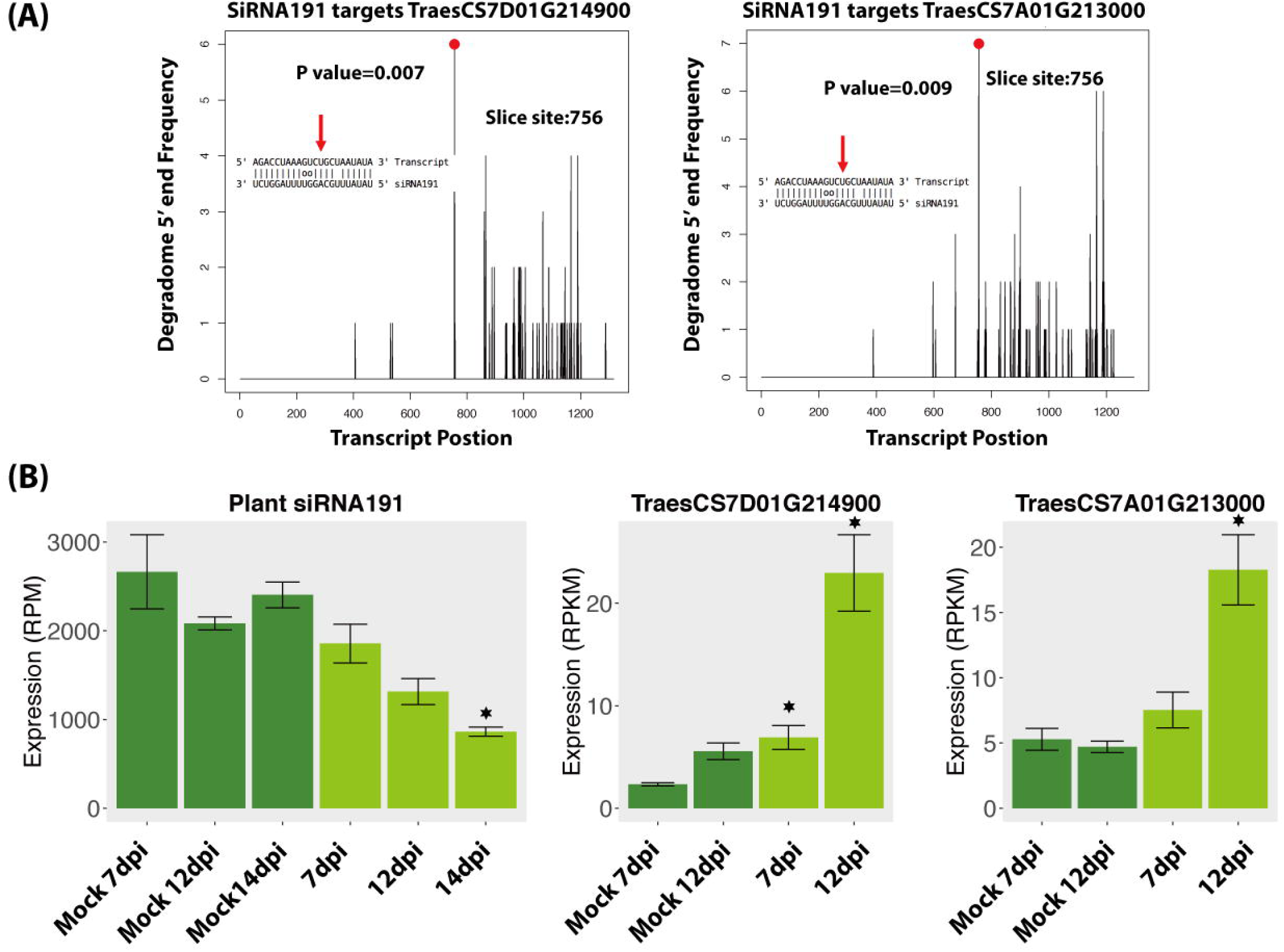
Wheat downregulates siRNA191 to upregulate two wheat RLK during the infections. (A) Degradome results of siRNA191 and its target genes in plant. (B) Expressions of siRNA191 and its target genes in plant. The asterisk indicates that the adjusted p value is under 0.05.

Conversely, 191 plant sRNAs were significantly upregulated in the infections, among which 39 were miRNAs (15 novel miRNAs) and 152 were siRNAs (Figure 5C, Table S6). Among these induced or extremely highly expressed (RPM above 100) plant sRNAs, 176 sRNAs (47 miRNAs and 129 siRNAs) were predicted to target and silence 690 wheat genes (Table S8). Interestingly, 20 (3%) of these candidates targeted genes were resistance genes and 77 (11.2%) were receptor like kinases (RLKs), which are considered to play central roles in plant immunity. Besides, 12 (2%) genes encode ABC transporters and 11 (1.5%) genes encode Auxin response factor which are also genes involved in the plant immune system (Figure 7A).

**Figure 7.**
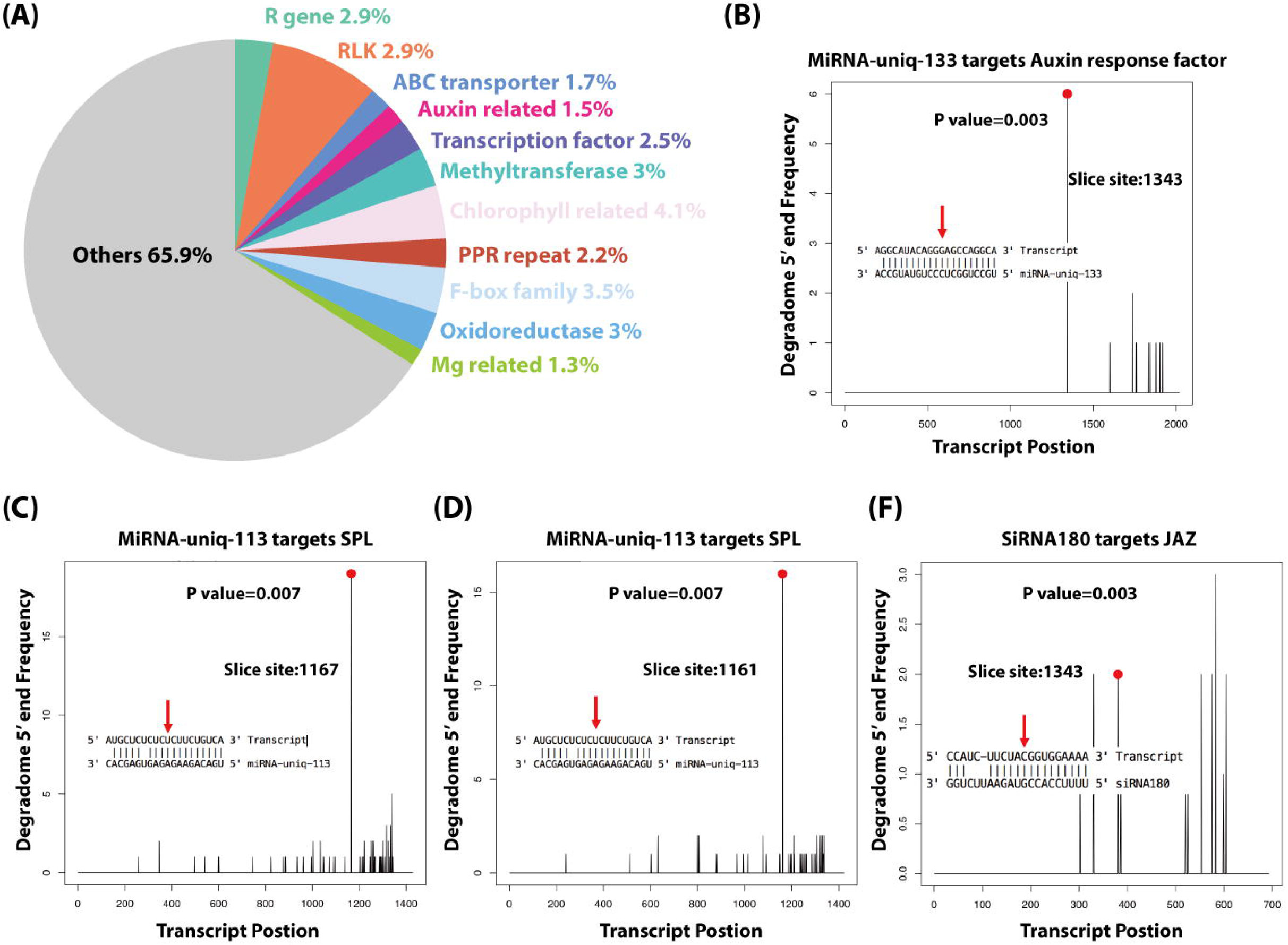
Wheat induces sRNAs to regulate wheat genes as an immune response against *Z. tritici*. (A) The functions of the wheat genes, which were downregulated and predicted to be targeted by wheat induced sRNAs. (B,C,D,E) Degradome analysis of the wheat genes targeted by miRNA-uniq-133, miRNA-uniq-113 and siRNA180.

We confirmed the cleavage of 5 transcripts among these 690 genes (Figure 7, S1). MiRNA-uniq-133 (miRNA160 family) was detected to silence TraesCS6A01G222300.1 which encodes an auxin response factor (Figure 7B). Two Squamosa promoter-binding-like gene (SPL) were negatively regulated by miRNA-uniq-113 (belonging to miRNA156 family), which was dramatically upregulated at 7dpi and 12dpi (Figure 7C,D). Jasmonate zim-domain (JAZ) protein, which negatively regulates Jasmonate (JA) signaling in plants, was silenced by siRNA180 at 7dpi (Figure 7E). SiRNA60 was also detected to target TraesCS3D01G339300.1, which encodes potassium transporter. But the low expression of siRNA60 and the high category of the degradome peak (in Category 3) suggests that it was not a real target (Figure S2). These findings suggest that wheat induced sRNAs to regulate plant gene expressions as a response to *Z. tritici* infections.

In plant-pathogen interactions, plants can as well export miRNAs into pathogen cells to influence pathogen virulence (Wang et al., 2016). We predicted 190 wheat siRNAs that were upregulated during the infections or extremely highly expressed (RPM above 100), and could target and downregulate 1115 fungal genes (Table S9). Interestingly, these targeted fungal genes were significantly enriched in “vesicle-mediated transport” GO term (Figure 8). We detected that two of these 1115 fungal genes could be cleaved by wheat sRNAs (Figure S3). One gene is g1915, which encodes an Antibiotic biosynthesis monooxygenase domain (ABM) protein. The other gene is g9791, which encodes FAD dependent oxidoreductase. These two fungal genes were only downregulated at the beginning of the disease cycle and recovered their expressions from 14dpi (Figure S3). However, it has to be noted that the degradome coverage over those genes was very low and the slice site of g1915 transcript belong to Category 3 (Figure S3), which suggests that is not a real target.

**Figure 8.**
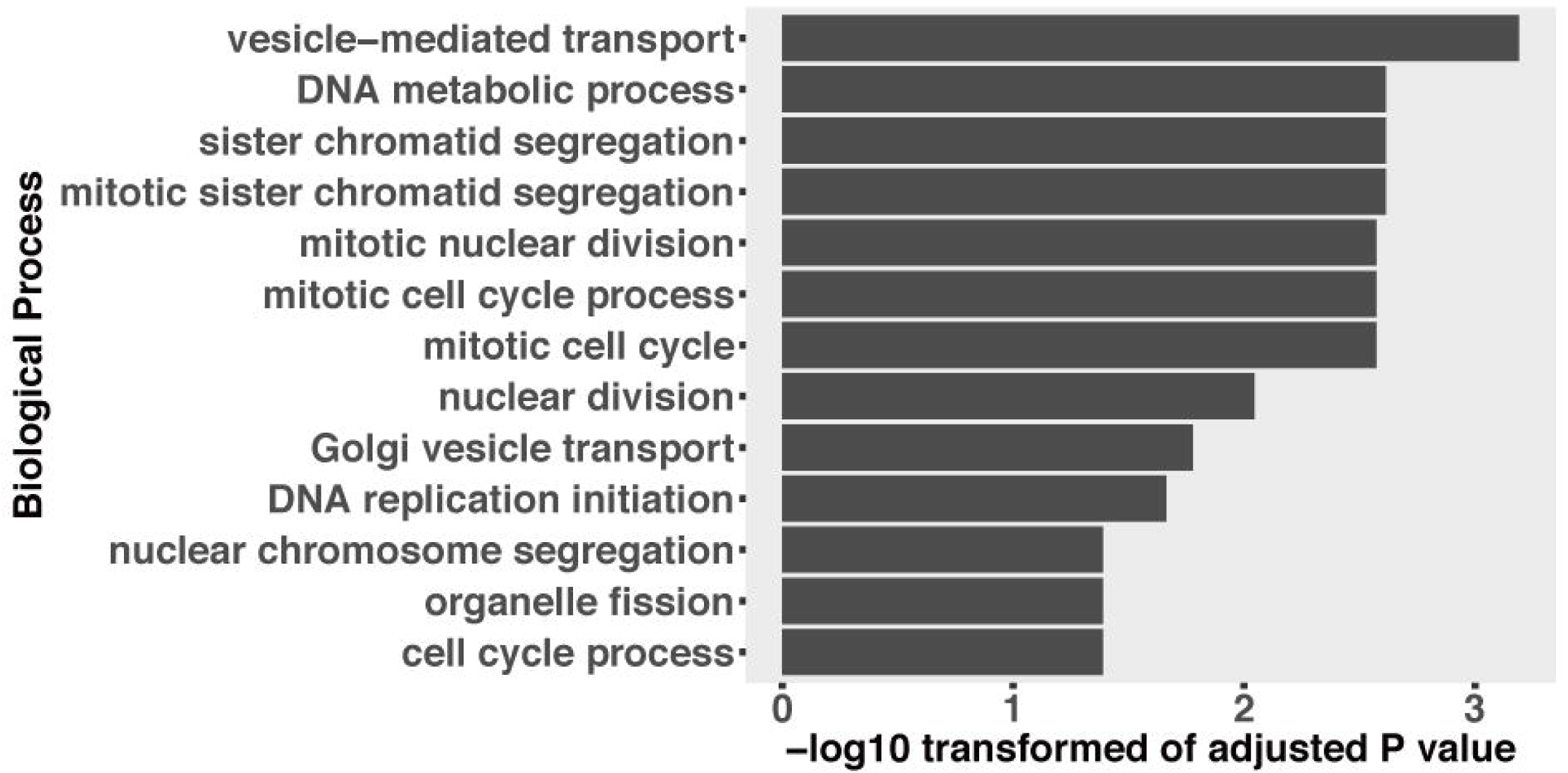
GO enrichment of fungal genes, which were downregulated and predicted to be targeted by induced plant sRNAs during the infection.

## Discussion

sRNAs have been shown to play important roles in plant-pathogen interactions, where RNAi is triggered naturally in a cross-kingdom manner (Nowara et al., 2010; Weiberg et al., 2013; Wang et al., 2016; Zhang et al., 2016; Wang et al., 2017; Cai et al., 2018). Here we combined sRNA-Seq, mRNA-Seq and degradome sequencing from samples covering key stages during the infection cycle to analyze the bidirectional RNAi in the wheat and *Z. tritici* pathosystem. We analyzed all fungal sRNAs and plant sRNAs in the infection samples, as well as in the fungus growing *in vitro* and non-infected plant mocks. By comparing the expression of sRNAs between infection and control samples, we found the infection induced sRNAs for both the fungus and the plant. The composition of these infection induced sRNAs changed during infections, suggesting that different sRNAs were induced or that they are loaded into different Ago proteins. By using degradome sequencing to validate the target predictions, we did not find a clear evidence that fungal sRNAs could cleave wheat defense related genes. However, our results suggest that wheat can regulate sRNAs in response to *Z. tritici* infections.

### RNAi does not play important roles in *Z. tritici* during wheat infection

Fungal sRNAs can hijack the plant host RNAi machinery to silence plant defense related genes and facilitate fungal infection (Weiberg et al., 2013; Wang et al., 2017). In our analysis, the predicted targets of the fungal sRNAs induced during wheat infections were significantly enriched in chlorophyll biosynthesis (Figure 3A). The inhibition of chlorophyll biosynthesis is consistent with the chlorotic symptoms observed from 12dpi. Chloroplasts are essential for the response to multiple environmental stimuli and for the biosynthesis of plant hormones (Shapiguzov et al., 2012). Besides, chloroplast can also produce reactive oxygen species (ROS) as an immune response to pathogens (Galvez-Valdivieso and Mullineaux, 2010). To successfully infect plants, pathogens can export effectors to target chloroplast to prevent chloroplastic ROS (Zabala et al., 2015). Here, we suggest that fungal sRNAs could target chlorophyll genes to disturb plant immunity. Also, the wheat genes targeted by the induced fungal sRNAs were also enriched in “microtubule-based movement” (Figure 3A). Plant microtubules need to be reorganized in response to symbiotic and pathogenic organisms to allow the proper intracellular trafficking activated in response to pathogen infection (Hardham, 2013). By disturbing the plant microtubules reorganization, may facilitate the fungal infection. Additionally, we found that plant RLKs were predicted to be targeted by fungal sRNAs (Table S3). These findings provide the possibility that the fungal sRNAs may be transported into plant cells and affect the plant immunity system. But all of these wheat genes mentioned above were not detected to be cleaved in the degradome analysis. We only detected one wheat gene that may be cleaved by fungal sRNAs. The wheat gene was TraesCS3B01G136500 and encodes a RSF-1 protein. (Figure 3D). But the expression of the corresponding fungal sRNA56 was very low (Figure 3E,F). Additionally, the frequency of the cleaved transcript from TraesCS3B01G136500 was very low. All these results suggest that this wheat gene was not a real target, at least in the canonical way.

We also predicted 39 fungal sRNA that could regulate fungal genes *in planta*. But none of these targeted fungal genes were detected to be cleaved in the degradome sequencing. The fungal Dicer gene was barely expressed in the infections, which leaded to a decrease of sRNA genesis by the fungus (Figure 4). This is supported by the lower expressions of sRNAs observed *in planta* than *in vitro*. In addition, deletion of the *Z. tritici* Dicer gene has been found to have no effect on fungal virulence, and it failed to block the disease (Kettles et al., 2018). These results, together with our findings, suggest minor roles of the *Z. tritici* Dicer for fungal virulence. Moreover, we did not find any strong evidence that fungal sRNAs could cleave plant transcripts. These data suggest that fungal sRNAs are not playing important roles in PTGS during the wheat-Z. *tritici* interaction. However, even if the *Z. tritici* sRNAs do not play a direct role in the interaction with wheat, they may play a role at specific developmental stages of the fungus, as it has been shown for other plant pathogenic fungi (Kim et al., 2015; Raman et al., 2017; Zeng et al., 2018). Also, we cannot discard the possibility that *Z. tritici* sRNAs may be acting at the transcriptional or translational level.

### Wheat sRNAs were induced in response to *Z. tritici*

Previous studies have reported that plants can employ sRNAs as a rapid defense response against pathogens. MiRNA393 is induced to target auxin related genes and repress auxin signaling in response to bacterial pathogens (Achard et al., 2007; Chen et al., 2007; Navarro et al., 2008; Grant and Jones, 2009). MYB transcription factors, which are involved largely in plant development and defense against abiotic and biotic stress, can also be targeted and regulated by the plant sRNAs (Samad et al., 2017). Additionally, plants also regulate the expressions of resistance genes by sRNAs under the pathogen attack (Zhai et al., 2011; Shivaprasad et al., 2012; Fei et al., 2013).

Here we found that 191 wheat sRNAs were downregulated during the fungal infections (Table S6). These downregulated wheat sRNAs were predicted to target R genes and RLKs in mocks (Table S7). Additionally, the cleavage of the transcripts from two RLKs were confirmed in the degradome analysis (Figure 6). These results suggest that during the infections, wheat could downregulate specific sRNAs triggering the upregulation of the target genes, especially the R genes and RLKs, and increase the plant defense ability in response to *Z. tritici* infections.

We also found that 191 wheat sRNAs were induced during the fungal infection (Figure 5C, Table S6). The composition of these induced sRNAs were quite different to the plant sRNAs expressed in the mocks (Figure 5B). Among these, 176 sRNAs (47miRNAs and 129 siRNAs) were predicted to target and silence 690 wheat genes (Table S8). These wheat genes included R genes, RLKs, ABC transporters and Auxin response factors, which are known to play central roles in the plant immunity (Figure 7A). These induced sRNA regulation of defense related genes suggests an immune response in wheat against *Z. tritici*. Besides, many transcription factor genes were also predicted to be targeted by these induced sRNAs, including SPL transcription factors, F-box domain proteins, MYB transcription factors and basic/helix-loop-helix (bHLH) proteins (Figure 7A). These results suggest that transcription factors can also be regulated by plant sRNAs as a response to the fungal pathogens. We confirmed the cleavage of five transcripts among these 690 wheat genes. They encode one auxin response factor, two SPL proteins, one JAZ protein and one potassium transporter (Figure 7). The induced sRNA mediated auxin related gene here is consistent with the response observed during plant infection by bacterial pathogens (Navarro et al, 2008). When plants are under biotic stress, sRNAs are induced to regulate transcription factors as an immune response (Tsuda and Somssich, 2015). Here we confirmed that SPL transcription factors are mainly involved in this sRNA mediated immune response. Besides, by silencing the JAZ gene with the induced sRNAs, the plant can trigger an enhance JA signaling as an immune response against the fungal pathogen. Thus, we show clear evidence that wheat will induce a group of specific sRNAs to regulate wheat genes as an immune response to *Z. tritici*.

In HIGS, plants can transport RNA modules and target pathogen genes to reduce pathogen virulence (Nowara et al., 2010; Baulcombe, 2015). It has been proved recently that plants use extracellular vesicles to send sRNAs into fungal cells (Cai et al., 2018). These finding suggest that the RNA molecules can be transported to fungal pathogens naturally. We predicted 1115 fungal genes that could be targeted by 190 wheat sRNAs. These wheat sRNAs were either upregulated or extremely highly expressed during the infections (Table S9). Interestingly, the targeted fungal genes were enriched in the “vesicle-mediated transport” GO term (Figure 8). Fungal Extracellular Vesicles (EVs) are essential to transport proteins, glycans pigments, nucleic acids and lipids (Rodrigues et al., 2015; Rodrigues and Casadevall, 2018). Fungal EVs are also responsible to pathogenesis during the infection (Rodrigues et al., 2007; Albuquerque et al., 2008; Rodrigues et al., 2008; Vallejo et al., 2011; Vargas et al., 2015), including mediating the transport of virulent effectors into plant cells (Rodrigues et al., 2008). A recent study also proved that plant miRNAs can be transported by EVs into fungal cells to affect fungal virulence (Cai et al., 2018). Here our results suggest that wheat sRNAs may also target fungal EVs related genes to interfere fungal virulence. However, we only detected two fungal genes that were cleaved by the wheat sRNAs (Figure S3). Fungal g1915, which encodes an ABM protein, involved in diverse biological processes, including metabolism, transcription, translation and biosynthesis of secondary metabolites was cleaved by plant siRNA180. The other one is fungal g9791, which encodes FAD dependent oxidoreductase, was targeted by plant miRNA-uniq-21. These two fungal genes play fundamental roles in fungal growth and they are downregulated at the beginning of the disease cycle. However, these two fungal genes were not completely silenced, and the expression of these two genes was recovered at 14dpi, when the fungus started the necrotrophic growth phase. But as we discussed before, the degradome results did not provide support for the cleavage of the transcripts from these two genes. These results suggest that wheat is not able to use sRNAs to silence fungal genes. These results are supported by the finding that dsRNAs generated from RNA virus vectors *in planta* are not effective in triggering gene silencing in *Z. tritici* during the fungal infection (Kettles et al., 2018).

In conclusion, our results suggest that there is no natural cross-kingdom RNAi by PTGS between these two interacting species. But we found that wheat can use sRNAs to regulate the plant defenses during *Z. tritici* infection. These findings contribute to improve our understanding of this pathosystem.

## Acknowledgements

SRNA and mRNA sequencing were performed at the Functional Genomics Center Zurich (FGCZ). The degradome sequencing was performed by Vertis Biotechnology AG (Germany). We thank Carolina S. Francisco for providing the RNA samples of the fungus *in vitro* used as control. We also thank Alexis Sarazin for his suggestions for the bioinformatics analysis, scientific discussions, and critical reading of the manuscript.

## Funding

Xin Ma is funded by a PSC-Syngenta PhD fellowship. Laboratory facilities were provided by the Genetic Diversity Center (GDC) of ETH Zurich.

## Authors’ contribution

X.M., N.B., and J.P.-G. conceived and designed experiments. X.M. was responsible for the bioinformatics analysis. X.M., N.B. and J.P.-G. wrote the paper.

## Supplementary information

**Figure S1. The expressions of wheat miRNA-uniq-133, miRNA-uniq-113, siRNA180 and siRNA60 and their target genes in wheat genome.**

**Figure S2. Degradome results of wheat siRNA60 and its target gene in plant.**

**Figure S3. Degradome result and expressions of two wheat genes and their target genes in fungi.**

**Table S1. Predicted sRNAs in *Z. tritici*.**

**Table S2. Differentially expressed fungal sRNAs between *in planta* and *in vitro* growth.**

**Table S3. Induced or highly expressed fungal sRNAs that target and downregulate wheat genes.**

**Table S4. Induced or highly expressed fungal sRNAs that target and downregulate fungal genes *in planta*.**

**Table S5. Predicted miRNAs and siRNAs in wheat.**

**Table S6. Differentially expressed wheat sRNAs.**

**Table S7. The downregulated wheat sRNAs and their target upregulated wheat genes in the infections.**

**Table S8. Induced or highly expressed wheat sRNAs that target and downregulate wheat genes in the infections.**

**Table S8. Induced or highly expressed wheat sRNAs that target and downregulate fungal genes in the infections.**

## Reference

Achard, P., Baghour, M., Chapple, A., Hedden, P., Van Der Straeten, D., Genschik, P., et al. (2007). The plant stress hormone ethylene controls floral transition via DELLA-dependent regulation of floral meristem-identity genes. Proc Natl Acad Sci U S A 104(15), 6484–6489. doi: 10.1073/pnas.0610717104.

Addo-Quaye, C., Snyder, J.A., Park, Y.B., Li, Y.F., Sunkar, R., and Axtell, M.J. (2009). Sliced microRNA targets and precise loop-first processing of MIR319 hairpins revealed by analysis of the Physcomitrella patens degradome. RNA 15(12), 2112–2121. doi: 10.1261/rna.1774909.

Albuquerque, P.C., Nakayasu, E.S., Rodrigues, M.L., Frases, S., Casadevall, A., Zancope-Oliveira, R.M., et al. (2008). Vesicular transport in Histoplasma capsulatum: an effective mechanism for trans-cell wall transfer of proteins and lipids in ascomycetes. Cell Microbiol 10(8), 1695–1710. doi: 10.1111/j.1462–5822.2008.01160.x.

Alexa, A., and Rahnenfuhrer, J. (2016). topGO: Enrichment Analysis for Gene Ontology. R package version 2.32.0.

Allen, E., Xie, Z., Gustafson, A.M., and Carrington, J.C. (2005). microRNA-directed phasing during trans-acting siRNA biogenesis in plants. Cell 121(2), 207–221. doi: 10.1016/j.cell.2005.04.004.

Anders, S., Pyl, P.T., and Huber, W. (2015). HTSeq--a Python framework to work with high-throughput sequencing data. Bioinformatics 31(2), 166–169. doi: 10.1093/bioinformatics/btu638.

Appels, R., Eversole, K., Feuillet, C., Keller, B., Rogers, J., Stein, N., et al. (2018). Shifting the limits in wheat research and breeding using a fully annotated reference genome. Science 361(6403). doi: 10.1126/science.aar7191.

Axtell, M.J. (2013). Classification and Comparison of Small RNAs from Plants. Annual Review of Plant Biology, Vol 64 64, 137–159. doi: 10.1146/annurev-arplant-050312-120043.

Axtell, M.J., and Meyers, B.C. (2018). Revisiting Criteria for Plant MicroRNA Annotation in the Era of Big Data. Plant Cell 30(2), 272–284. doi: 10.1105/tpc.17.00851.

Baulcombe, D.C. (2015). VIGS, HIGS and FIGS: small RNA silencing in the interactions of viruses or filamentous organisms with their plant hosts. Curr Opin Plant Biol 26, 141–146. doi: 10.1016/j.pbi.2015.06.007.

Bolger, A.M., Lohse, M., and Usadel, B. (2014). Trimmomatic: a flexible trimmer for Illumina sequence data. Bioinformatics 30(15), 2114–2120. doi: 10.1093/bioinformatics/btu170.

Bologna, N.G., and Voinnet, O. (2014). The diversity, biogenesis, and activities of endogenous silencing small RNAs in Arabidopsis. Annu Rev Plant Biol 65, 473–503. doi: 10.1146/annurev-arplant-050213-035728.

Borges, F., and Martienssen, R.A. (2015). The expanding world of small RNAs in plants. Nat Rev Mol Cell Biol 16(12), 727–741. doi: 10.1038/nrm4085.

Brousse, C., Liu, Q., Beauclair, L., Deremetz, A., Axtell, M.J., and Bouche, N. (2014). A non-canonical plant microRNA target site. Nucleic Acids Res 42(8), 5270–5279. doi: 10.1093/nar/gku157.

Cai, Q., Qiao, L., Wang, M., He, B., Lin, F.M., Palmquist, J., et al. (2018). Plants send small RNAs in extracellular vesicles to fungal pathogen to silence virulence genes. Science. doi: 10.1126/science.aar4142.

Camacho, C., Coulouris, G., Avagyan, V., Ma, N., Papadopoulos, J., Bealer, K., et al. (2009). BLAST plus : architecture and applications. Bmc Bioinformatics 10. doi: Artn 421 10.1186/1471-2105-10-421.

Carthew, R.W., and Sontheimer, E.J. (2009). Origins and Mechanisms of miRNAs and siRNAs. Cell 136(4), 642–655. doi: 10.1016/j.cell.2009.01.035.

Chang, S.S., Zhang, Z.Y., and Liu, Y. (2012). RNA Interference Pathways in Fungi: Mechanisms and Functions. Annual Review of Microbiology, Vol 66 66, 305–323. doi: 10.1146/annurev-micro-092611-150138.

Chen, Z., Agnew, J.L., Cohen, J.D., He, P., Shan, L., Sheen, J., et al. (2007). Pseudomonas syringae type III effector AvrRpt2 alters Arabidopsis thaliana auxin physiology. Proc Natl Acad Sci U S A 104(50), 20131–20136. doi: 10.1073/pnas.0704901104.

Creasey, K.M., Zhai, J., Borges, F., Van Ex, F., Regulski, M., Meyers, B.C., et al. (2014). miRNAs trigger widespread epigenetically activated siRNAs from transposons in Arabidopsis. Nature 508(7496), 411–415. doi: 10.1038/nature13069.

Dai, X., Zhuang, Z., and Zhao, P.X. (2018). psRNATarget: a plant small RNA target analysis server (2017 release). Nucleic Acids Res 46(W1), W49–W54. doi: 10.1093/nar/gky316.

Fei, Q., Xia, R., and Meyers, B.C. (2013). Phased, secondary, small interfering RNAs in posttranscriptional regulatory networks. Plant Cell 25(7), 2400–2415. doi: 10.1105/tpc.113.114652.

Felippes, F.F., and Weigel, D. (2009). Triggering the formation of tasiRNAs in Arabidopsis thaliana: the role of microRNA miR173. EMBO Rep 10(3), 264–270. doi: 10.1038/embor.2008.247.

Francisco, C.S., Ma, X., Zwyssig, M.M., McDonald, B.A., and Palma-Guerrero, J. (2018). Coping with stress: morphological changes in response to environmental stimuli in a fungal plant pathogen. bioRxiv. doi: 10.1101/372078.

Galvez-Valdivieso, G., and Mullineaux, P.M. (2010). The role of reactive oxygen species in signalling from chloroplasts to the nucleus. Physiologia Plantarum 138(4), 430–439. doi: 10.1111/j.1399-3054.2009.01331.x.

Goodwin, S.B., M’Barek S, B., Dhillon, B., Wittenberg, A.H., Crane, C.F., Hane, J.K., et al. (2011). Finished genome of the fungal wheat pathogen Mycosphaerella graminicola reveals dispensome structure, chromosome plasticity, and stealth pathogenesis. PLoS Genet 7(6), e1002070. doi: 10.1371/journal.pgen.1002070.

Gordon, A., and Hannon, G. (2010). "Fastx-toolkit. FASTQ/A short-reads preprocessing tools (unpublished)".).

Grant, M.R., and Jones, J.D. (2009). Hormone (dis)harmony moulds plant health and disease. Science 324(5928), 750–752. doi: 10.1126/science.1173771.

Griffiths-Jones, S. (2004). The microRNA Registry. Nucleic Acids Res 32(Database issue), D109–111. doi: 10.1093/nar/gkh023.

Griffiths-Jones, S., Grocock, R.J., van Dongen, S., Bateman, A., and Enright, A.J. (2006). miRBase: microRNA sequences, targets and gene nomenclature. Nucleic Acids Res 34(Database issue), D140–144. doi: 10.1093/nar/gkj112.

Griffiths-Jones, S., Saini, H.K., van Dongen, S., and Enright, A.J. (2008). miRBase: tools for microRNA genomics. Nucleic Acids Res 36(Database issue), D154–158. doi: 10.1093/nar/gkm952.

Guleria, P., Mahajan, M., Bhardwaj, J., and Yadav, S.K. (2011). Plant small RNAs: biogenesis, mode of action and their roles in abiotic stresses. Genomics Proteomics Bioinformatics 9(6), 183–199. doi: 10.1016/S1672-0229(11)60022-3.

Hardham, A.R. (2013). Microtubules and biotic interactions. Plant Journal 75(2), 278–289. doi: 10.1111/tpj.12171.

Hardwick, N.V., Jones, D.R., and Slough, J.E. (2001). Factors affecting diseases of winter wheat in England and Wales, 1989–98. Plant Pathology 50(4), 453–462. doi: DOI 10.1046/j.1365-3059.2001.00596.x.

Hutvagner, G., and Simard, M.J. (2008). Argonaute proteins: key players in RNA silencing. Nat Rev Mol Cell Biol 9(1), 22–32. doi: 10.1038/nrm2321.

Johnson, N.R., Yeoh, J.M., Coruh, C., and Axtell, M.J. (2016). Improved Placement of Multi-mapping Small RNAs. G3 (Bethesda) 6(7), 2103–2111. doi: 10.1534/g3.116.030452.

Kamthan, A., Chaudhuri, A., Kamthan, M., and Datta, A. (2015). Small RNAs in plants: recent development and application for crop improvement. Front Plant Sci 6, 208. doi: 10.3389/fpls.2015.00208.

Kersey, P.J., Allen, J.E., Allot, A., Barba, M., Boddu, S., Bolt, B.J., et al. (2018). Ensembl Genomes 2018: an integrated omics infrastructure for non-vertebrate species. Nucleic Acids Res 46(D1), D802–D808. doi: 10.1093/nar/gkx1011.

Kettles, G.J., Hofinge, B.J., Hu, P., Bayon, C., Rudd, J.J., Balmer, D., et al. (2018). Analysis of small RNA silencing in Zymoseptoria tritici – wheat interactions. submitted.

Kim, D., Pertea, G., Trapnell, C., Pimentel, H., Kelley, R., and Salzberg, S.L. (2013). TopHat2: accurate alignment of transcriptomes in the presence of insertions, deletions and gene fusions. Genome Biol 14(4), R36. doi: 10.1186/gb-2013-14-4-r36.

Kim, H.K., Jo, S.M., Kim, G.Y., Kim, D.W., Kim, Y.K., and Yun, S.H. (2015). A Large-Scale Functional Analysis of Putative Target Genes of Mating-Type Loci Provides Insight into the Regulation of Sexual Development of the Cereal Pathogen Fusarium graminearum. Plos Genetics 11(9). doi: ARTN e1005486 10.1371/journal.pgen.1005486.

Kozomara, A., and Griffiths-Jones, S. (2011). miRBase: integrating microRNA annotation and deep-sequencing data. Nucleic Acids Res 39(Database issue), D152–157. doi: 10.1093/nar/gkq1027.

Kozomara, A., and Griffiths-Jones, S. (2014). miRBase: annotating high confidence microRNAs using deep sequencing data. Nucleic Acids Res 42(Database issue), D68–73. doi: 10.1093/nar/gkt1181.

Langmead, B., Trapnell, C., Pop, M., and Salzberg, S.L. (2009). Ultrafast and memory-efficient alignment of short DNA sequences to the human genome. Genome Biol 10(3), R25. doi: 10.1186/gb-2009-10-3-r25.

Lee, A.H., Hurley, B., Felsensteiner, C., Yea, C., Ckurshumova, W., Bartetzko, V., et al. (2012). A bacterial acetyltransferase destroys plant microtubule networks and blocks secretion. PLoS Pathog 8(2), e1002523. doi: 10.1371/journal.ppat.1002523.

Lee, H.C., Chang, S.S., Choudhary, S., Aalto, A.P., Maiti, M., Bamford, D.H., et al. (2009). qiRNA is a new type of small interfering RNA induced by DNA damage. Nature 459(7244), 274–U163. doi: 10.1038/nature08041.

Lee, H.C., Li, L., Gu, W., Xue, Z., Crosthwaite, S.K., Pertsemlidis, A., et al. (2010). Diverse pathways generate microRNA-like RNAs and Dicer-independent small interfering RNAs in fungi. Mol Cell 38(6), 803–814. doi: 10.1016/j.molcel.2010.04.005.

Li, F., Pignatta, D., Bendix, C., Brunkard, J.O., Cohn, M.M., Tung, J., et al. (2012). MicroRNA regulation of plant innate immune receptors. Proceedings of the National Academy of Sciences of the United States of America 109(5), 1790–1795. doi: 10.1073/pnas.1118282109.

Liu, J.D., Carmell, M.A., Rivas, F.V., Marsden, C.G., Thomson, J.M., Song, J.J., et al. (2004). Argonaute2 is the catalytic engine of mammalian RNAi. Science 305(5689), 1437–1441. doi: 10.1126/science.1102513.

Ma, X., Keller, B., McDonald, B.A., Palma-Guerrero, J., and Wicker, T. (2018). Comparative transcriptomics reveals how wheat responds to infection by Zymoseptoria tritici. Molecular Plant-Microbe Interactions, MPMI-10-17-0245-R.

Marshall, R., Kombrink, A., Motteram, J., Loza-Reyes, E., Lucas, J., Hammond-Kosack, K.E., et al. (2011). Analysis of two in planta expressed LysM effector homologs from the fungus Mycosphaerella graminicola reveals novel functional properties and varying contributions to virulence on wheat. Plant Physiol 156(2), 756–769. doi: 10.1104/pp.111.176347.

Matzke, M.A., and Mosher, R.A. (2014). RNA-directed DNA methylation: an epigenetic pathway of increasing complexity. Nat Rev Genet 15(6), 394–408. doi: 10.1038/nrg3683.

Menzies, I.J., Youard, L.W., Lord, J.M., Carpenter, K.L., van Klink, J.W., Perry, N.B., et al. (2016). Leaf colour polymorphisms: a balance between plant defence and photosynthesis. Journal of Ecology 104(1), 104–113. doi: 10.1111/1365-2745.12494.

Mi, S.J., Cai, T., Hu, Y.G., Chen, Y., Hodges, E., Ni, F.R., et al. (2008). Sorting of small RNAs into Arabidopsis argonaute complexes is directed by the 5 ' terminal nucleotide. Cell 133(1), 116–127. doi: 10.1016/j.cell.2008.02.034.

Moazed, D. (2009). Small RNAs in transcriptional gene silencing and genome defence. Nature 457(7228), 413–420. doi: 10.1038/nature07756.

Navarro, L., Dunoyer, P., Jay, F., Arnold, B., Dharmasiri, N., Estelle, M., et al. (2006). A plant miRNA contributes to antibacterial resistance by repressing auxin signaling. Science 312(5772), 436–439. doi: 10.1126/science.1126088.

Navarro, L., Jay, F., Nomura, K., He, S.Y., and Voinnet, O. (2008). Suppression of the microRNA pathway by bacterial effector proteins. Science 321(5891), 964–967. doi: 10.1126/science.1159505.

Nicolas, F.E., and Ruiz-Vazquez, R.M. (2013). Functional Diversity of RNAi-Associated sRNAs in Fungi. International Journal of Molecular Sciences 14(8), 15348–15360. doi: 10.3390/ijms140815348.

Nolan, T., Braccini, L., Azzalin, G., De Toni, A., Macino, G., and Cogoni, C. (2005). The post-transcriptional gene silencing machinery functions independently of DNA methylation to repress a LINE1-like retrotransposon in Neurospora crassa. Nucleic Acids Res 33(5), 1564–1573. doi: 10.1093/nar/gki300.

Nowara, D., Gay, A., Lacomme, C., Shaw, J., Ridout, C., Douchkov, D., et al. (2010). HIGS: host-induced gene silencing in the obligate biotrophic fungal pathogen Blumeria graminis. Plant Cell 22(9), 3130–3141. doi: 10.1105/tpc.110.077040.

O’Driscoll, A., Kildea, S., Doohan, F., Spink, J., and Mullins, E. (2014). The wheat-Septoria conflict: a new front opening up? Trends in Plant Science 19(9), 602–610. doi: 10.1016/j.tplants.2014.04.011.

Palma-Guerrero, J., Ma, X., Torriani, S.F., Zala, M., Francisco, C.S., Hartmann, F.E., et al. (2017). Comparative Transcriptome Analyses in Zymoseptoria tritici Reveal Significant Differences in Gene Expression Among Strains During Plant Infection. Mol Plant Microbe Interact 30(3), 231–244. doi: 10.1094/MPMI-07-16-0146-R.

Park, G.G., Park, J.J., Yoon, J., Yu, S.N., and An, G. (2010). A RING finger E3 ligase gene, Oryza sativa Delayed Seed Germination 1 (OsDSG1), controls seed germination and stress responses in rice. Plant Mol Biol 74(4–5), 467–478. doi: 10.1007/s11103-010-9687-3.

Plissonneau, C., Hartmann, F.E., and Croll, D. (2018). Pangenome analyses of the wheat pathogen Zymoseptoria tritici reveal the structural basis of a highly plastic eukaryotic genome. BMC Biol 16(1), 5. doi: 10.1186/s12915-017-0457-4.

Plissonneau, C., Sturchler, A., and Croll, D. (2016). The Evolution of Orphan Regions in Genomes of a Fungal Pathogen of Wheat. MBio 7(5). doi: 10.1128/mBio.01231-16.

Rajagopalan, R., Vaucheret, H., Trejo, J., and Bartel, D.P. (2006). A diverse and evolutionarily fluid set of microRNAs in Arabidopsis thaliana. Genes Dev 20(24), 3407–3425. doi: 10.1101/gad.1476406.

Raman, V., Simon, S.A., Demirci, F., Nakano, M., Meyers, B.C., and Donofrio, N.M. (2017). Small RNA Functions Are Required for Growth and Development of Magnaporthe oryzae. Mol Plant Microbe Interact 30(7), 517–530. doi: 10.1094/MPMI-11-16-0236-R.

Robinson, M.D., McCarthy, D.J., and Smyth, G.K. (2010). edgeR: a Bioconductor package for differential expression analysis of digital gene expression data. Bioinformatics 26(1), 139–140. doi: 10.1093/bioinformatics/btp616.

Robinson, M.D., and Oshlack, A. (2010). A scaling normalization method for differential expression analysis of RNA-seq data. Genome Biol 11(3), R25. doi: 10.1186/gb-2010-11-3-r25.

Rodrigues, M.L., and Casadevall, A. (2018). A two-way road: novel roles for fungal extracellular vesicles. Mol Microbiol 110(1), 11–15. doi: 10.1111/mmi.14095.

Rodrigues, M.L., Godinho, R.M.C., Zamith-Miranda, D., and Nimrichter, L. (2015). Traveling into Outer Space: Unanswered Questions about Fungal Extracellular Vesicles. Plos Pathogens 11(12). doi: ARTN e1005240 10.1371/journal.ppat.1005240.

Rodrigues, M.L., Nimrichter, L., Oliveira, D.L., Frases, S., Miranda, K., Zaragoza, O., et al. (2007). Vesicular polysaccharide export in Cryptococcus neoformans is a eukaryotic solution to the problem of fungal trans-cell wall transport. Eukaryotic Cell 6(1), 48–59. doi: 10.1128/Ec.00318-06.

Rodrigues, M.L., Nimrichter, L., Oliveira, D.L., Nosanchuk, J.D., and Casadevall, A. (2008). Vesicular Trans-Cell Wall Transport in Fungi: A Mechanism for the Delivery of Virulence-Associated Macromolecules? Lipid Insights 2, 27–40.

Romano, N., and Macino, G. (1992). Quelling: transient inactivation of gene expression in Neurospora crassa by transformation with homologous sequences. Mol Microbiol 6(22), 3343–3353.

Rudd, J.J., Kanyuka, K., Hassani-Pak, K., Derbyshire, M., Andongabo, A., Devonshire, J., et al. (2015). Transcriptome and metabolite profiling of the infection cycle of Zymoseptoria tritici on wheat reveals a biphasic interaction with plant immunity involving differential pathogen chromosomal contributions and a variation on the hemibiotrophic lifestyle definition. Plant Physiol 167(3), 1158–1185. doi: 10.1104/pp.114.255927.

Ruiz-Ferrer, V., and Voinnet, O. (2009). Roles of plant small RNAs in biotic stress responses. Annu Rev Plant Biol 60, 485–510. doi: 10.1146/annurev.arplant.043008.092111.

Samad, A.F.A., Sajad, M., Nazaruddin, N., Fauzi, I.A., Murad, A.M.A., Zainal, Z., et al. (2017). MicroRNA and Transcription Factor: Key Players in Plant Regulatory Network. Front Plant Sci 8, 565. doi: 10.3389/fpls.2017.00565.

Sanchez-Vallet, A., McDonald, M.C., Solomon, P.S., and McDonald, B.A. (2015). Is Zymoseptoria tritici a hemibiotroph? Fungal Genet Biol 79, 29–32. doi: 10.1016/j.fgb.2015.04.001.

Shahid, S., and Axtell, M.J. (2014). Identification and annotation of small RNA genes using ShortStack. Methods 67(1), 20–27. doi: 10.1016/j.ymeth.2013.10.004.

Shapiguzov, A., Vainonen, J.P., Wrzaczek, M., and Kangasjarvi, J. (2012). ROS-talk - how the apoplast, the chloroplast, and the nucleus get the message through. Frontiers in Plant Science 3. doi: ARTN 292 10.3389/fpls.2012.00292.

Shivaprasad, P.V., Chen, H.M., Patel, K., Bond, D.M., Santos, B.A., and Baulcombe, D.C. (2012). A microRNA superfamily regulates nucleotide binding site-leucine-rich repeats and other mRNAs. Plant Cell 24(3), 859–874. doi: 10.1105/tpc.111.095380.

Siomi, M.C., Sato, K., Pezic, D., and Aravin, A.A. (2011). PIWI-interacting small RNAs: the vanguard of genome defence. Nat Rev Mol Cell Biol 12(4), 246–258. doi: 10.1038/nrm3089.

Sunkar, R., Chinnusamy, V., Zhu, J.H., and Zhu, J.K. (2007). Small RNAs as big players in plant abiotic stress responses and nutrient deprivation. Trends in Plant Science 12(7), 301–309. doi: 10.1016/j.tplants.2007.05.001.

Sunkar, R., and Zhu, J.K. (2004). Novel and stress-regulated microRNAs and other small RNAs from Arabidopsis. Plant Cell 16(8), 2001–2019. doi: 10.1105/tpc.104.022830.

Torres-Martinez, S., and Ruiz-Vazquez, R.M. (2017). The RNAi Universe in Fungi: A Varied Landscape of Small RNAs and Biological Functions. Annual Review of Microbiology, Vol 71 71, 371–391. doi: 10.1146/annurev-micro-090816-093352.

Tsuda, K., and Somssich, I.E. (2015). Transcriptional networks in plant immunity. New Phytol 206(3), 932–947. doi: 10.1111/nph.13286.

Vagin, V.V., Sigova, A., Li, C., Seitz, H., Gvozdev, V., and Zamore, P.D. (2006). A distinct small RNA pathway silences selfish genetic elements in the germline. Science 313(5785), 320–324. doi: 10.1126/science.1129333.

Vallejo, M.C., Matsuo, A.L., Ganiko, L., Medeiros, L.C., Miranda, K., Silva, L.S., et al. (2011). The pathogenic fungus Paracoccidioides brasiliensis exports extracellular vesicles containing highly immunogenic alpha-Galactosyl epitopes. Eukaryot Cell 10(3), 343–351. doi: 10.1128/EC.00227-10.

Vargas, G., Rocha, J.D.B., Oliveira, D.L., Albuquerque, P.C., Frases, S., Santos, S.S., et al. (2015). Compositional and immunobiological analyses of extracellular vesicles released by Candida albicans. Cellular Microbiology 17(3), 389–407. doi: 10.1111/cmi.12374.

Wang, B., Sun, Y., Song, N., Zhao, M., Liu, R., Feng, H., et al. (2017). Puccinia striiformis f. sp. tritici microRNA-like RNA 1 (Pst-milR1), an important pathogenicity factor of Pst, impairs wheat resistance to Pst by suppressing the wheat pathogenesis-related 2 gene. New Phytol 215(1), 338–350. doi: 10.1111/nph.14577.

Wang, M., Weiberg, A., Lin, F.M., Thomma, B.P., Huang, H.D., and Jin, H. (2016). Bidirectional cross-kingdom RNAi and fungal uptake of external RNAs confer plant protection. Nat Plants 2, 16151. doi: 10.1038/nplants.2016.151.

Weiberg, A., Wang, M., Bellinger, M., and Jin, H. (2014). Small RNAs: a new paradigm in plant-microbe interactions. Annu Rev Phytopathol 52, 495–516. doi: 10.1146/annurev-phyto-102313-045933.

Weiberg, A., Wang, M., Lin, F.M., Zhao, H., Zhang, Z., Kaloshian, I., et al. (2013). Fungal small RNAs suppress plant immunity by hijacking host RNA interference pathways. Science 342(6154), 118–123. doi: 10.1126/science.1239705.

Yang, X., and Li, L. (2011). miRDeep-P: a computational tool for analyzing the microRNA transcriptome in plants. Bioinformatics 27(18), 2614–2615. doi: 10.1093/bioinformatics/btr430.

Zabala, M.D.T., Littlejohn, G., Jayaraman, S., Studholme, D., Bailey, T., Lawson, T., et al. (2015). Chloroplasts play a central role in plant defence and are targeted by pathogen effectors. Nature Plants 1(6). doi: Artn 15074 10.1038/Nplants.2015.74.

Zeng, W.P., Wang, J., Wang, Y., Lin, J., Fu, Y.P., Xie, J.T., et al. (2018). Dicer-Like Proteins Regulate Sexual Development via the Biogenesis of Perithecium-Specific MicroRNAs in a Plant Pathogenic Fungus Fusarium graminearum. Frontiers in Microbiology 9. doi: ARTN 818 10.3389/fmicb.2018.00818.

Zhai, J.X., Jeong, D.H., De Paoli, E., Park, S., Rosen, B.D., Li, Y.P., et al. (2011). MicroRNAs as master regulators of the plant NB-LRR defense gene family via the production of phased, trans-acting siRNAs. Genes & Development 25(23), 2540–2553. doi: 10.1101/gad.177527.111.

Zhang, T., Zhao, Y.L., Zhao, J.H., Wang, S., Jin, Y., Chen, Z.Q., et al. (2016). Cotton plants export microRNAs to inhibit virulence gene expression in a fungal pathogen. Nat Plants 2(10), 16153. doi: 10.1038/nplants.2016.153.

Zhang, W., Gao, S., Zhou, X., Chellappan, P., Chen, Z., Zhou, X., et al. (2011). Bacteria-responsive microRNAs regulate plant innate immunity by modulating plant hormone networks. Plant Mol Biol 75(1–2), 93–105. doi: 10.1007/s11103-010-9710-8.

