## Supplementary figures and images for "Small RNA bidirectional crosstalk during the interaction between wheat and *Zymoseptoria tritici*" (from another version of this preprint: version 1, 2018-12-19)

### Figure S1

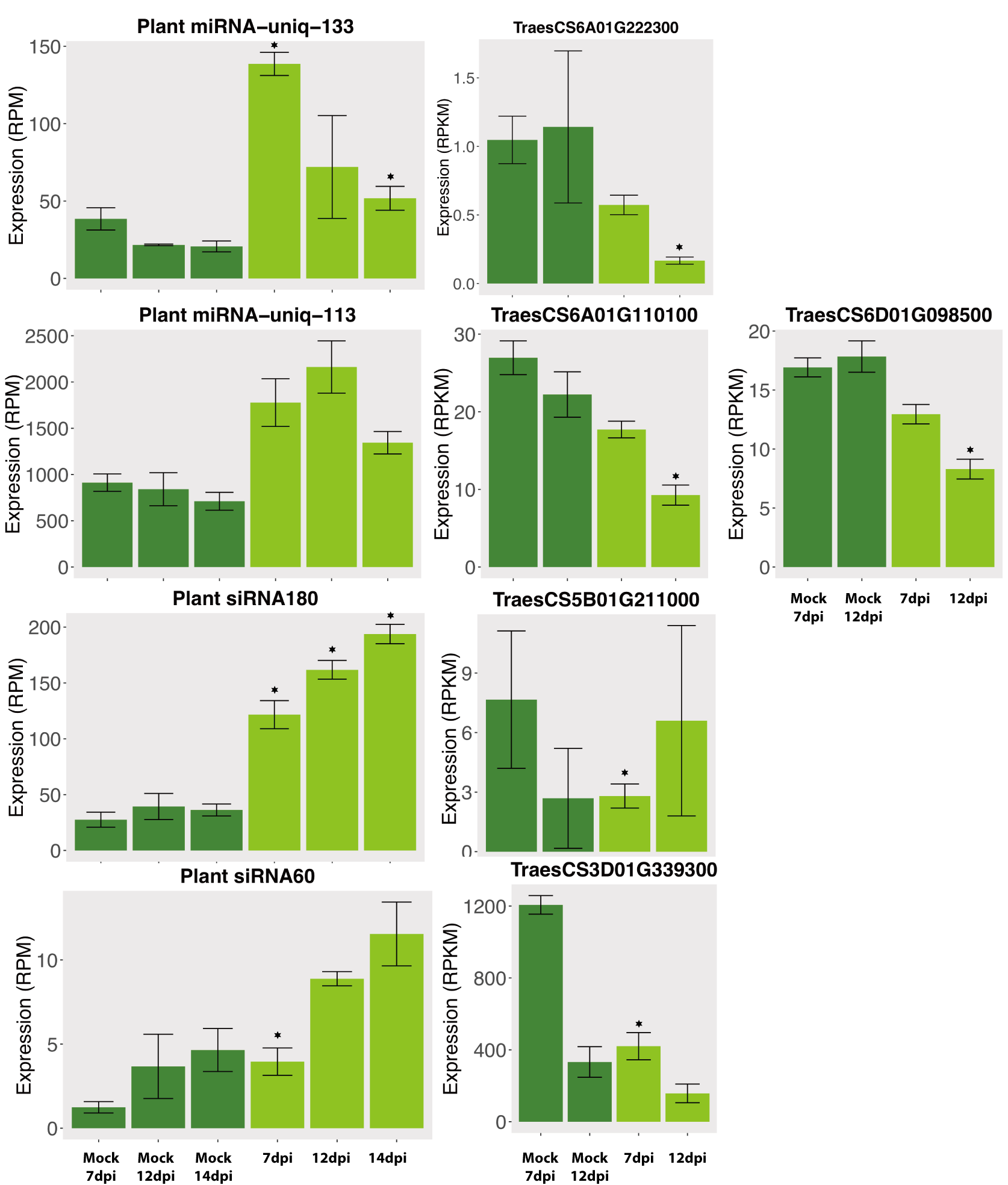

### Figure S2

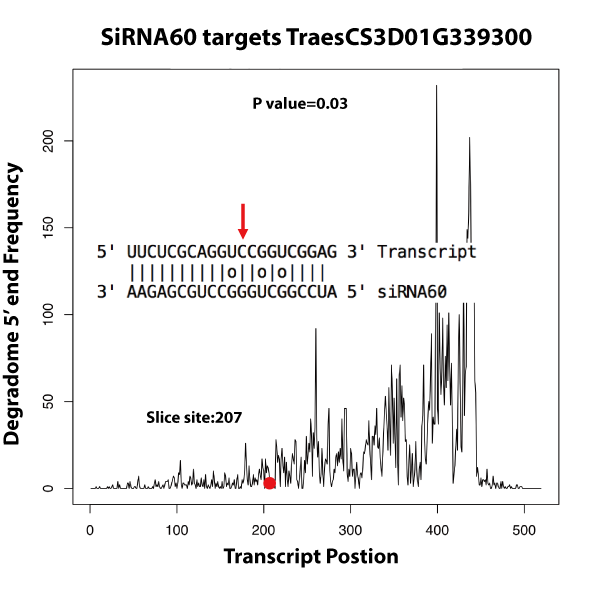

### Figure S3

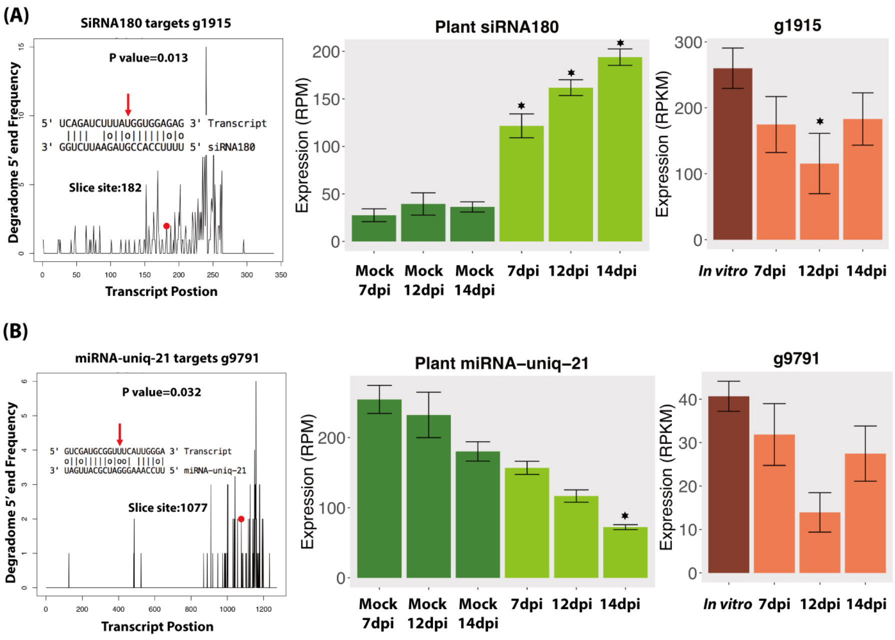
